# Light activates the translational regulatory GCN2 kinase via reactive oxygen species emanating from the chloroplast

**DOI:** 10.1101/794362

**Authors:** Ansul Lokdarshi, Ju Guan, Ricardo A Urquidi Camacho, Sung Ki Cho, Philip Morgan, Madison Leonard, Masaki Shimono, Brad Day, Albrecht G von Arnim

## Abstract

Cytosolic mRNA translation is subject to global and mRNA-specific controls. Phosphorylation of translation initiation factor eIF2α anchors a reversible switch that represses translation globally. The stress-responsive GCN2 kinase is the only known kinase for eIF2α in *Arabidopsis*. Here we show that conditions that generate reactive oxygen species (ROS) in the chloroplast, such as dark-light transitions, high light, and the herbicide methyl viologen all rapidly activated the GCN2 kinase, whereas mitochondrial and ER stress did not. In addition, GCN2 activation was light dependent and mitigated by photosynthesis inhibitors and ROS quenchers. Accordingly, seedling growth of multiple *gcn2* mutant alleles was retarded under conditions of excess light, implicating the GCN2-eIF2α pathway in responses to light and associated ROS. Once activated, the GCN2 kinase preferentially suppressed the ribosome loading of mRNAs for functions such as mitochondrial ATP synthesis, the chloroplast thylakoids, vesicle trafficking, and translation. The transcriptome of *gcn2* mutants was sensitized to abiotic stress, including oxidative stress, as well as innate immune responses. Accordingly, *gcn2* displayed defects in immune priming by the fungal elicitor, chitin. In conclusion, we provide evidence that reactive oxygen species produced by the photosynthetic apparatus help to activate the highly conserved GCN2 kinase, leading to eIF2α phosphorylation and thus affecting the status of the cytosolic protein synthesis apparatus.

## INTRODUCTION

Reactive oxygen species (ROS) are constantly produced as byproducts of plant cellular metabolism and also serve as versatile secondary messengers (Foyer and Noctor, 2016; Mignolet-Spruyt et al., 2016; Choudhury et al., 2017; Mittler, 2017; Mullineaux et al., 2018; Waszczak et al., 2018). Labelled as being a double-edged sword, plants tightly regulate the ROS balance between production and scavenging by various enzymatic and non-enzymatic mechanisms (Das and Roychoudhury, 2014; Foyer and Noctor, 2016; Mittler, 2017). As signaling molecules, these mechanisms function in a variety of physiological and developmental programs, including root development (Tsukagoshi, 2016), the pathogen-induced hypersensitive response (Camejo et al., 2016), and stomatal closure (Pei et al., 2000; Ehonen et al., 2018). These normal programs are quite often perturbed by a variety of external biotic and abiotic cues, which make ROS a cytotoxic agent. The resulting adverse effects among others include oxidative damage to nucleic acids, carbohydrates, and especially to lipids and proteins (Jacques et al., 2013; Demidchik, 2015).

Being both beneficial and deleterious, plant ROS production is carefully compartmentalized to the apoplast, endoplasmic reticulum, chloroplasts, mitochondria and peroxisomes (Tripathy and Oelmuller, 2012; Del Rio and Lopez-Huertas, 2016; Czarnocka and Karpinski, 2018). Under conditions of active photosynthesis, chloroplasts serve as the major producers of ROS (Schmitt et al., 2014). Arabidopsis plants adapted to low light conditions and exposed to sudden high light intensities experience photo-oxidative stress resulting from the excess excitation energy (Mateo et al., 2004; Munoz and Munne-Bosch, 2018), a phenomenon whereby the amount of absorbed light energy exceeds the photosynthetic capacity (Li et al., 2009). Under excess light, singlet oxygen (^1^O_2_) and superoxide anion (O_2_^-1^) are overproduced at photosystems II and I respectively, which in turn result in higher levels of hydrogen peroxide (H_2_O_2_) (Asada, 2006; Mubarakshina et al., 2010; Dietz et al., 2016). When high light elevates organellar ROS production beyond the capacity for their detoxification, this triggers stress adaptive reprogramming by retrograde signaling to the nucleus (Dietz, 2015; Foyer and Noctor, 2016; Mignolet-Spruyt et al., 2016; Crisp et al., 2017).

Most of the work to dissect the role of ROS on gene expression has focused on transcriptional control (Vanderauwera et al., 2005; Alboresi et al., 2011; Lai et al., 2012; Vaahtera et al., 2014; Bode et al., 2016; Xu et al., 2017), which is inherently slow. Being rapidly reversible, regulation at the level of translation may regulate gene expression in a nimbler manner in response to rapidly changing light and ROS levels. Light is well known to affect cytosolic translation and generally has a stimulatory effect (Tang et al., 2003; Juntawong and Bailey-Serres, 2012; Liu et al., 2012; Missra et al., 2015; Moore et al., 2016; Merchante et al., 2017). The effect of ROS on cytosolic translation (Khandal et al., 2009) has remained understudied. The underlying mechanisms may involve redox-sensitive translation factors but are generally not well understood (Moore et al., 2016).

In mammals and yeast, translational control in response to a diverse range of stresses converges on the phosphorylation of translation initiation factor eIF2α (eIF2α-P), a regulatory process carried out by a family of up to four different kinases. Upon phosphorylation, eIF2α binds to eIF2B and inhibits its guanine nucleotide exchange activity, leading to a depletion of the ternary complex (eIF2-GTP-tRNA(i)Met), and ultimately, a decline in translation initiation and protein synthesis (Wek, 2018). In plants, the only known kinase for eIF2α is GCN2, which responds to diverse stress stimuli including inhibitors of amino acid and purine biosynthesis, wounding, UV light, hormones and bacterial infection by phosphorylating eIF2α (Akbudak et al., 2006; Lageix et al., 2008; Zhang et al., 2008; Luna et al., 2014; Sesma et al., 2017; Izquierdo et al., 2018; Liu et al., 2019). GCN2 is activated through the binding of various uncharged tRNAs, which accumulate during amino acid starvation and other stress conditions (Dong et al., 2000; Li et al., 2013; Anda et al., 2017).

In this study, we show that the GCN2 kinase is activated rapidly under excess light stress and H_2_O_2_-mediated oxidative stress. We conclude that this increase in GCN2 kinase activity and eIF2α-P is specific to a yet unknown signal emanating from the chloroplast because it can be controlled by the application of various plastidic redox modulators. Moreover, eIF2α phosphorylation in response to herbicide stress is strictly light-dependent. Seedlings with mutations in *GCN2* grow more slowly under continuous or high light stress. They are also defective in the global reorganization of the translatome that occurs after GCN2 is activated by herbicide. Interestingly, comparative transcriptome analysis between wild type and *gcn2* plants under herbicide stress reveals significant changes in mRNAs involved in the response to pathogens as well as abiotic stresses, including ROS metabolism. Taken together, the study presented herein shows how ROS emanating from the photosynthetic machinery in the chloroplast feed back to the cytosolic protein synthesis apparatus to balance overall energy and metabolic resources with demand.

## RESULTS

### GCN2 kinase activity is activated by excess light

The activity of the GCN2 kinase can be determined by monitoring the phosphorylation status of its primary substrate, the eukaryotic translation initiation factor eIF2α. GCN2 is known to be activated by a number of environmental signals, including herbicides that inhibit amino acid biosynthesis, UV light, and salicylic acid (Lageix et al., 2008; Zhang et al., 2008). To first determine if GCN2 is activated by light, 24-hr dark-adapted seedlings were exposed to white light (80 µEin m^-2^s^-1^). After dark-adaptation, the eIF2α-P level declined to the level in unstimulated plants, whereas white light induced intense eIF2α phosphorylation within 2 hr in dark adapted wild-type plants, but not in the *gcn2* mutant (**Fig. 1A, B**), indicating that the GCN2 kinase is responsive to light. GCN2 activity was fluence-rate dependent, and was rapidly elevated (30 min) by a moderately high light intensity (200 µEin m^-2^s^-1^) (**Fig. 1C**). In contrast, heat shock (i.e., 37 °C) did not trigger an increase in GCN2 activity albeit suppressed GCN2 basal activity under light (**Supplemental Figure 1**). Under normal long-day light-dark cycle conditions (i.e., 80 µEin m^-2^s^-1^, no dark adaptation), eIF2α-P rose to maximal levels during the light period, then declined early during the dark period (**Fig. 1D, E**). These results indicate that the GCN2 kinase is activated by white light and that responses to other signals should be cognizant of the diel dynamics of eIF2α-P. Once phosphorylated under high light (780 µEin m^-2^s^-1^), eIF2α-P remained elevated for over 72 hr (**Supplemental Figure 2**) indicating little adaptation to the high light condition. In contrast, the light-triggered eIF2α-P dissipated within 6 hr after shifting plants to darkness (**Supplemental Figure 3**).

**Figure 1.**
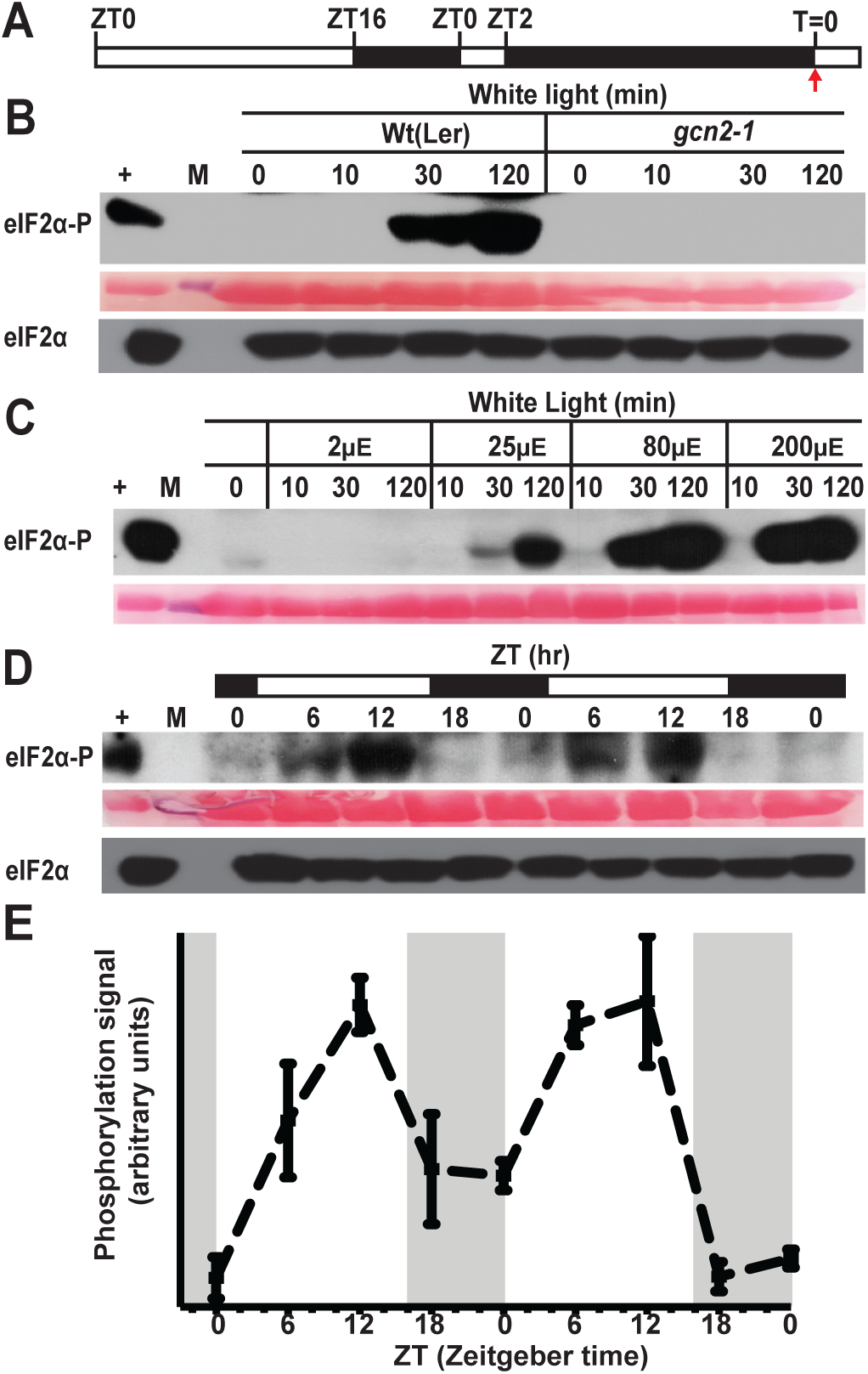
Excess light triggers GCN2-dependent eIF2α phosphorylation in a dose dependent manner. **(A)** Schematic of the light regimen. Seedlings were grown in a 16 hr light 8 hr dark cycle, followed by a 24 hr dark acclimation starting at zeitgeber time (ZT)2. The red arrow at T=0 indicates the beginning of excess light treatment and the start of sampling. **(B)** Immunoblot showing the time course of eIF2α phosphorylation in 14-days-old wild-type Landsberg (Wt (Ler)) and *gcn2-1* mutant (*gcn2-1*) seedlings subjected to excess light stress (White light) as described in panel (A). Upper panel: Probed with phospho-specific antibody against eIF2α-P (38kDa). A partially cropped band at the top represents nonspecific binding of the antibody. Middle panel: Rubisco large subunit (∼ 55kDa) as a loading control after Ponceau S staining of the blot. Lower panel: Probed with antibody against eIF2α (38kDa). (+), arbitrary amount of total protein extract from glyphosate treated Wt seedlings indicating unphosphorylated (eIF2α) or phosphorylated (eIF2α-P) protein; (10, 30, 120) sampling time in minutes; (M) Molecular weight marker. **(C)** Fluence rate dependence of eIF2α phosphorylation in Wt seedlings with 2, 25, 80 and 200 µEin m^-2^s^-1^ (µE) white light after dark acclimation as described in panel (A, B). **(D)** Diel time course of eIF2α phosphorylation over 48 hr in 14-days-old Wt seedlings grown under a 16 hr light (80 µEin m^-2^s^-1^) and 8 hr dark cycle. **(E)** Quantification of the diel time course of the eIF2α phosphorylation signal from immunoblots as shown in (D). Dark periods are shaded gray. Error bars represent standard error of the mean of three independent biological replicates.

**Figure 2.**
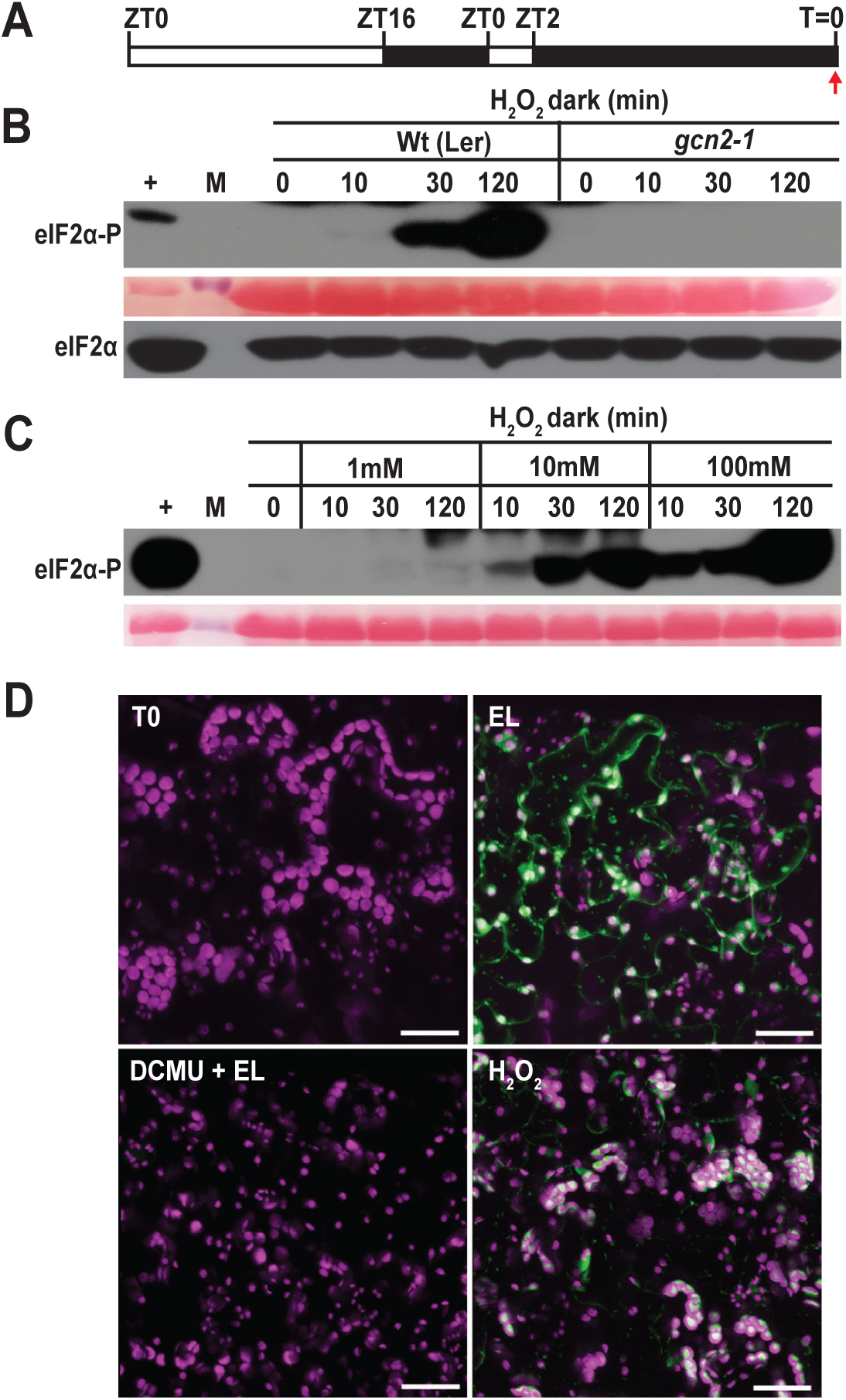
Effect of hydrogen peroxide on GCN2 activity, and microscopic examination of ROS in response to photosynthetic inhibitors. **(A)** Schematic of the light regimen. Seedlings were grown in a 16 hr light 8 hr dark cycle, dark-acclimated for 24 hr, and then sprayed with H_2_O_2_ in dark starting at T=0 (red arrow). **(B)** Time course of eIF2α phosphorylation in 14-days-old wild-type Landsberg (Wt(Ler)) and *gcn2-1* mutant (*gcn2-1*) treated with 10mM H_2_O_2_ for 0, 10, 30 and 120 minutes as described in panel (A). For details see legend to Fig. 1. **(C)** Treatment of Wt seedlings with 1, 10 and 100mM H_2_O_2_ after 24 hr dark acclimation as described in panel (A). **(D)** Representative images of 14-days-old Wt leaves stained with the ROS sensitive dye H_2_DCFDA after 24 hr dark acclimation as in (A). T0, dark acclimated control; EL, exposed to 80 µEin m^-2^s^-1^ white light for 30 minutes. DCMU+EL, treated with 8µM 3-(3,4-dichlorophenyl)-1,1-dimethylurea (DCMU) 30 minutes prior to exposure to 80 µEin m^-2^s^-1^ light. H_2_ O_2_, treated with 10mM H_2_O_2_ in the dark for 30 minutes. Oxidized H_2_DCFDA is represented in green and chlorophyll in magenta, and the superimposed view yields a white color. Scale bars are 25µm.

**Figure 3.**
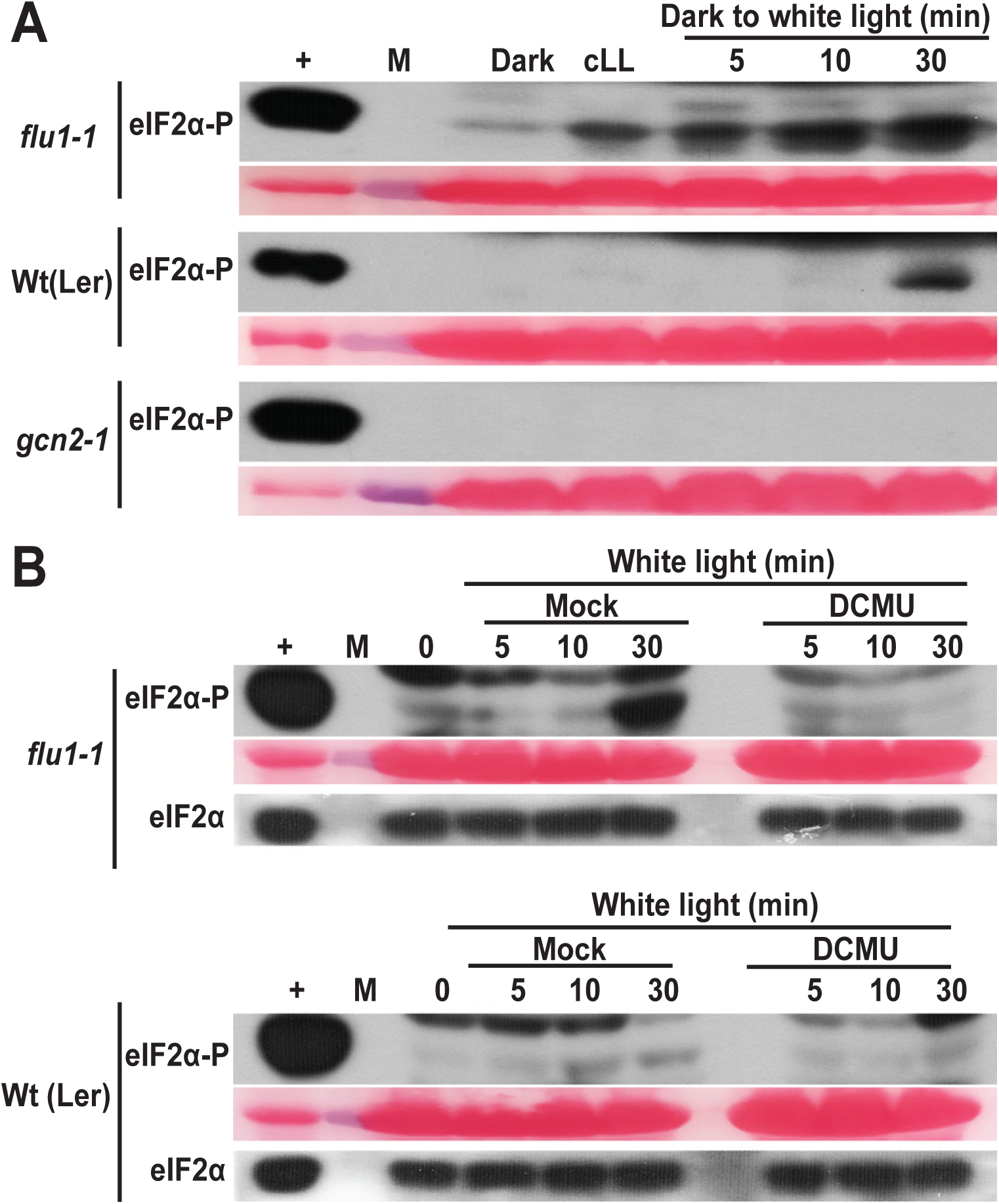
*flu* mutants show accelerated or elevated GCN2 activation by photosynthetic ROS. **(A)** Time course of eIF2α phosphorylation in rosette stage leaves of wild-type (Wt (Ler)) and *flu1-1* mutants. Plants were grown under continuous light (cLL), dark acclimated for 16 hr (Dark) and then re-exposed to light at 80 µEin m^-2^s^-1^ (Dark to white light). Note that eIF2α-P is elevated within 5 minutes in the *flu1-1* mutant. *gcn2-1* is included as a negative control. **(B)** eIF2α phosphorylation in *flu1-1* mutant seedlings and Wt (Ler) control after 12 days in continuous light, followed by 24 hr dark acclimation and re-exposure to light (80 µEin m^-2^s^-1^). Seedlings were sprayed with either DMSO (Mock), or 8µM DCMU, 30 minutes prior to light exposure. Time=0 was sampled right before the light treatment. For details see legend to Fig.1.

### GCN2 is activated by reactive oxygen species

Because excess light leads to a rapid accumulation of ROS (Asada, 1999; Asada, 2006; Galvez-Valdivieso et al., 2009) such as hydrogen peroxide, we tested whether GCN2 could be activated solely by H_2_O_2_. Indeed, ectopic H_2_O_2_ treatments under dark resulted in a dose-dependent activation of GCN2 as early as 10 min and this induction was reduced, although still detectable at doses as low as 1mM (**Fig. 2A, B, C**). In keeping with the hypothesis that excess light induces ROS accumulation, exposing dark-adapted seedlings to light triggered ROS accumulation in seedling leaves, as shown *in situ* with the ROS-sensitive dye 2’,7’-dichlorofluorescein diacetate (H_2_DCFDA) (**Fig. 2D**).

To gain closer insights about the identity of the ROS (H_2_O_2_, ^1^O_2_, O_2_^-1^) that can activate GCN2, eIF2α phosphorylation was analyzed with specific ROS inducers. The conditional *fluorescent* (*flu*) mutant generates singlet oxygen (^1^O_2_) in plastids within 1min upon a shift from prolonged darkness to light and therefore must be grown in continuous light for survival (Meskauskiene et al., 2001; op den Camp et al., 2003). Indeed, upon dark to light shift *flu1-1* mutant plants showed more rapid (<5min) eIF2α phosphorylation than wild-type, and the basal level of phosphorylation was elevated as well (**Fig. 3A**). Interestingly, the hypersensitive activation of GCN2 in the *flu1-1* mutant was suppressed in seedlings pretreated with the plastoquinone oxidizer, 3-(3,4-dichlorophenyl)-1,1-dimethyl urea (DCMU) **(Fig. 3B)**. ^1^O_2_ is rapidly converted to H_2_O_2_ in a non-enzymatic reaction utilizing either plastoquinone (PQ) or plastoquinol (PQH_2_) (Asada, 1999; Mubarakshina and Ivanov, 2010; Khorobrykh et al., 2015; Vetoshkina et al., 2017). Taken together, chloroplast singlet oxygen can serve as the trigger for the GCN2 kinase, notwithstanding that singlet oxygen may act via hydrogen peroxide.

### Herbicides rely on photosynthetic H_2_O_2_ to activate GCN2

GCN2 is activated by uncharged tRNA (Wek et al., 1995), including in plants (Li et al., 2013). In keeping with this, treatments with herbicides that inhibit amino acid biosynthesis activate GCN2 kinase (Lageix et al., 2008; Zhang et al., 2008). We tested three compounds that inhibit different branch points in amino acid biosynthesis; chlorosulfuron for branched-chain amino acids, glyphosate for aromatic amino acids, and glufosinate for glutamine synthetase. Surprisingly, all three inhibitors required light to activate GCN2 kinase (**Fig. 4A-D**). These data support the hypothesis that GCN2 cannot be activated solely by herbicides raising the levels of uncharged tRNA, yet is activated by a signal that requires light. We posit that ROS remains a candidate for this signal because treatment with a low concentration of chlorosulfuron herbicide led to ROS accumulation in the leaf (**Fig. 4E**), as does exposure to glyphosate and glufosinate (Faus et al., 2015; Takano et al., 2019). Moreover, the antioxidant ascorbate could delay the activation of GCN2 kinase by excess light (**Supplemental Figure 4)**.

**Figure 4.**
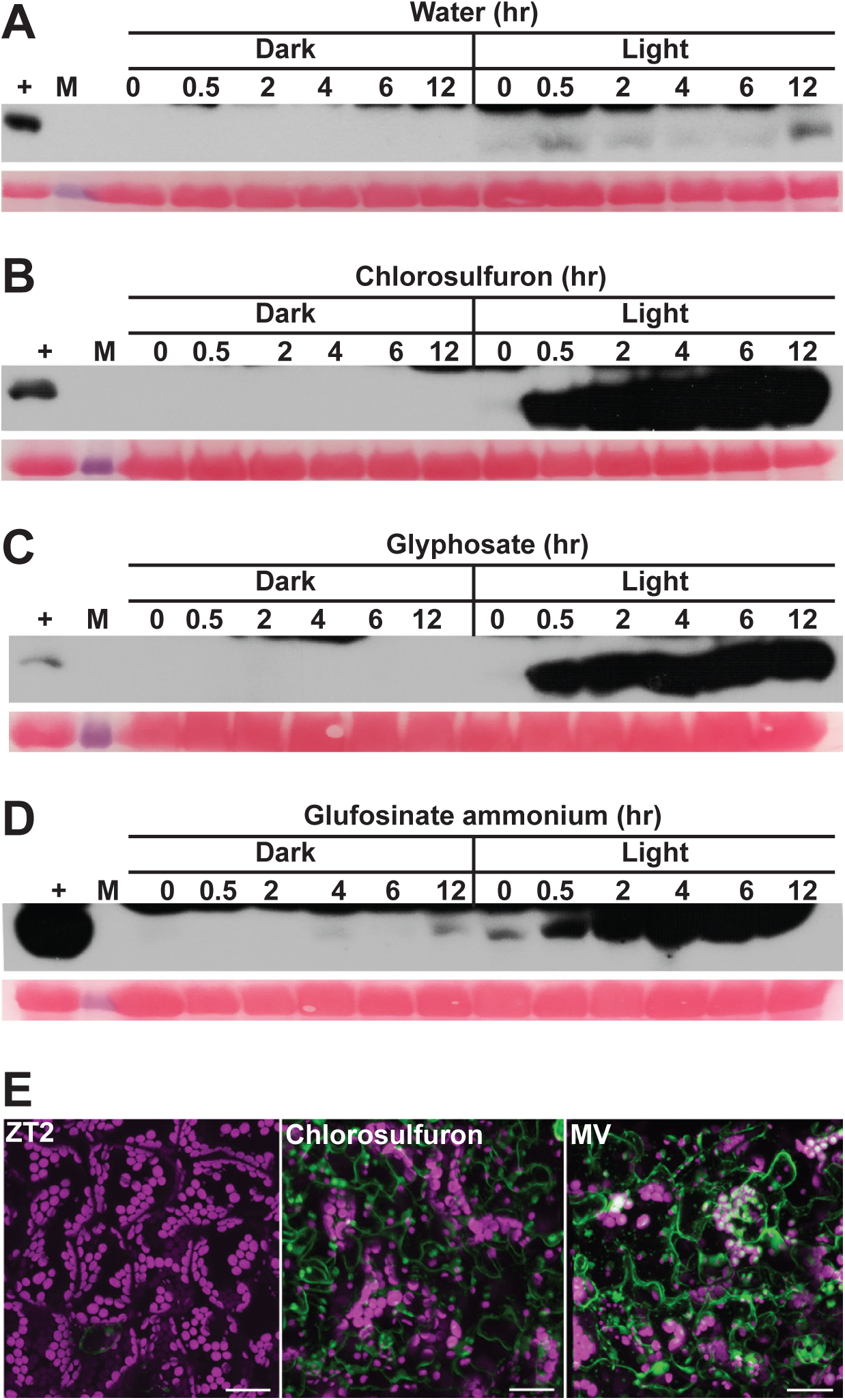
GCN2 activation with herbicide requires light. **(A-D)** Time course of eIF2α phosphorylation in 14-days-old wild-type Landsberg seedlings treated with **(A)** water only, **(B)** 0.6µM chlorosulfuron, **(C)** 150µM glyphosate, **(D)** 15µg/ml glufosinate ammonium under light and dark conditions. For treatments under dark and details for blots, see legend to Fig 1. **(E)** Chlorosulfuron triggers ROS accumulation under light as visualized with H_2_DCFDA. 14-days-old Wt leaves were sampled at ZT2 (left panel), then treated with 0.6µM chlorosulfuron for 1 hr starting at ZT2 (Chlorosulfuron). MV, treated with 20µM methyl viologen (MV) for 30 minutes. For details see legend to Fig 2D. Scale bars are 25µm.

To test the hypothesis that chlorosulfuron may act on GCN2 by imbalancing chloroplast ROS metabolism, wild-type seedlings were pretreated with photosynthetic inhibitors that manipulate the plastoquinone/plastoquinole (PQ/PQH_2_) pool before treatment with chlorosulfuron. Specifically, application of the plastoquinone oxidizer, (DCMU), or the reducer, 2,5-dibromo-3-methyl-6-isopropyl-p-benzoquinone (DBMIB) (Mateo et al., 2004; Kruk and Karpinski, 2006) both suppressed the effect of a near-saturating dose of chlorosulfuron (**Fig. 5A, B**). Taken together, these findings suggest that uncharged tRNAs cannot be the only activation signal for GCN2. A second signal appears to be needed as well, and given that ROS can induce GCN2 on its own, it is a candidate for this signal.

**Figure 5.**
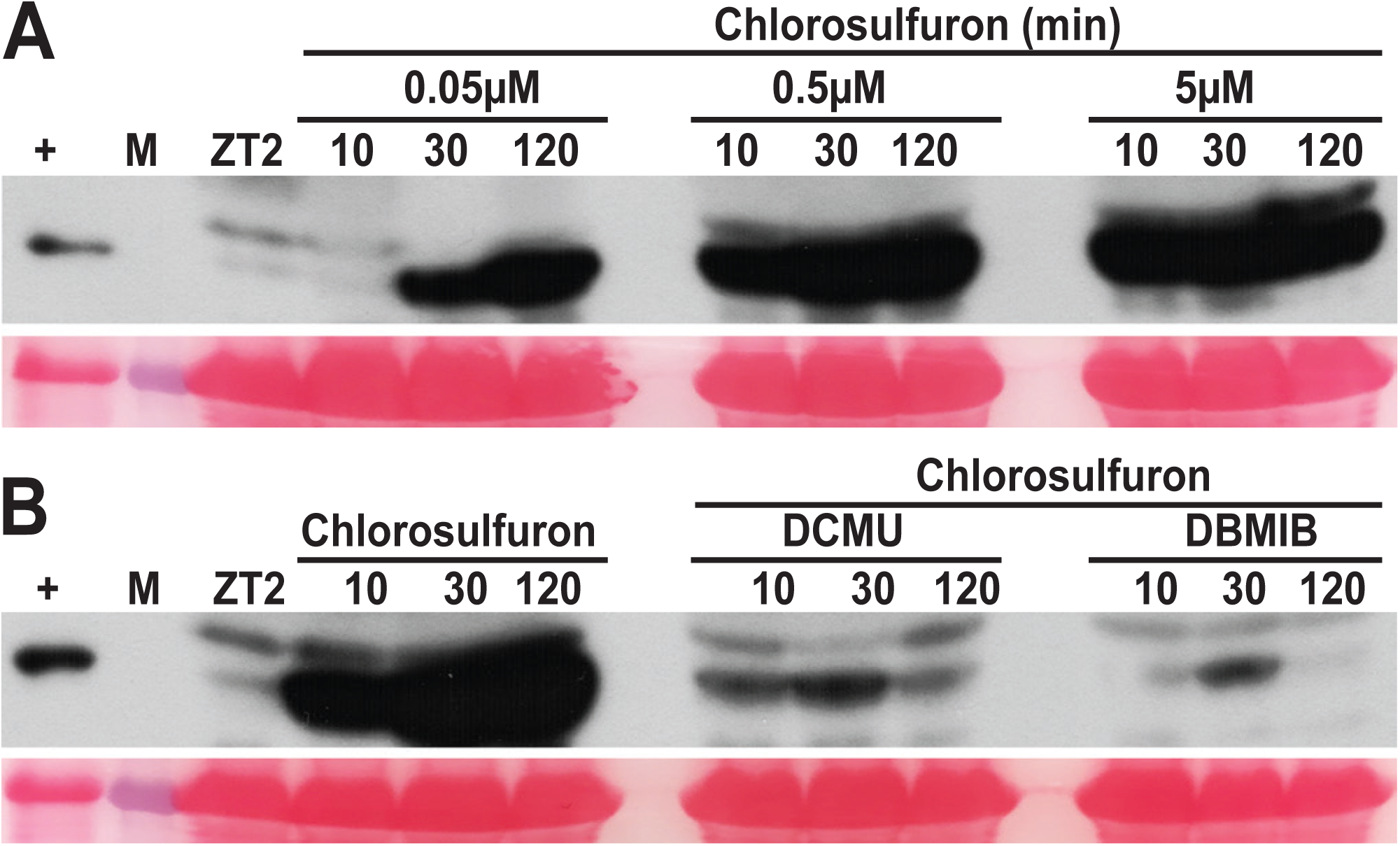
Photosynthetic inhibitors attenuate chlorosulfuron-triggered GCN2 activation. eIF2α phosphorylation in long day grown 14-days-old wild-type Landsberg seedlings. Treatments with herbicides started at ZT2 in the light. **(A)** Dose response for chlorosulfuron over a time course of up to 120 minutes. **(B)** eIF2α phosphorylation triggered by chlorosulfuron (0.5µM) was suppressed by pretreatment with 8µM of DCMU, or 16µM of 2,5-Dibromo-6-isopropyl-3-methyl-1,4-benzoquinone (DBMIB). DCMU and DBMIB were sprayed 30 minutes prior to chlorosulfuron treatment. For details see legend to Fig.1

### GCN2 supports seedling growth under conditions of excess light

The eIF2α phosphorylation data, above, support a role for GCN2 under high light. Under normal laboratory conditions, *gcn2-1* (Landsberg ecotype) mutants display few, if any, phenotypic abnormalities (Lageix et al., 2008; Zhang et al., 2008; Faus et al., 2015; Liu et al., 2015). After exposure to 3 days of continuous high light, the growth of *gcn2-1* mutant seedlings was retarded as compared to wild type, specifically in roots and by overall fresh weight (**Fig. 6A-C**), whereas seedlings grown in a regular day-night cycle were normal. Limited by the availability of only a single mutant allele (*gcn2-1*) in the Landsberg ecotype, we characterized two independent *gcn2* alleles in the Columbia (Col-0) ecotype, *gcn2-2* (SALK_032196) and *gcn2-3* (SALK_129334.2) (**Supplemental Figure 5 A-D**). The *gcn2-2* and *gcn2-3* alleles also had reduced root growth after 3 days of continuous high light (cHL, **Supplemental Figure 6A-B and 7**) and, following recovery in normal light, lower fresh weight in *gcn2-2* (**Supplemental Figure 6C**). However, other than *gcn2-1*, these Columbia alleles also had shorter roots under continuous normal light intensity (+3 days cLL, **Supplemental Figure 6 and 7**) suggesting that the sensitivity to a lesion in *GCN2* is ecotype-dependent. Taken together, given that three independent *gcn2* loss of function alleles had similar whole-seedling phenotypes, we conclude that GCN2 kinase has a physiological role in adaptation to excess light. However, other environmental conditions yet to be examined may well reveal additional roles for the GCN2 kinase.

**Figure 6.**
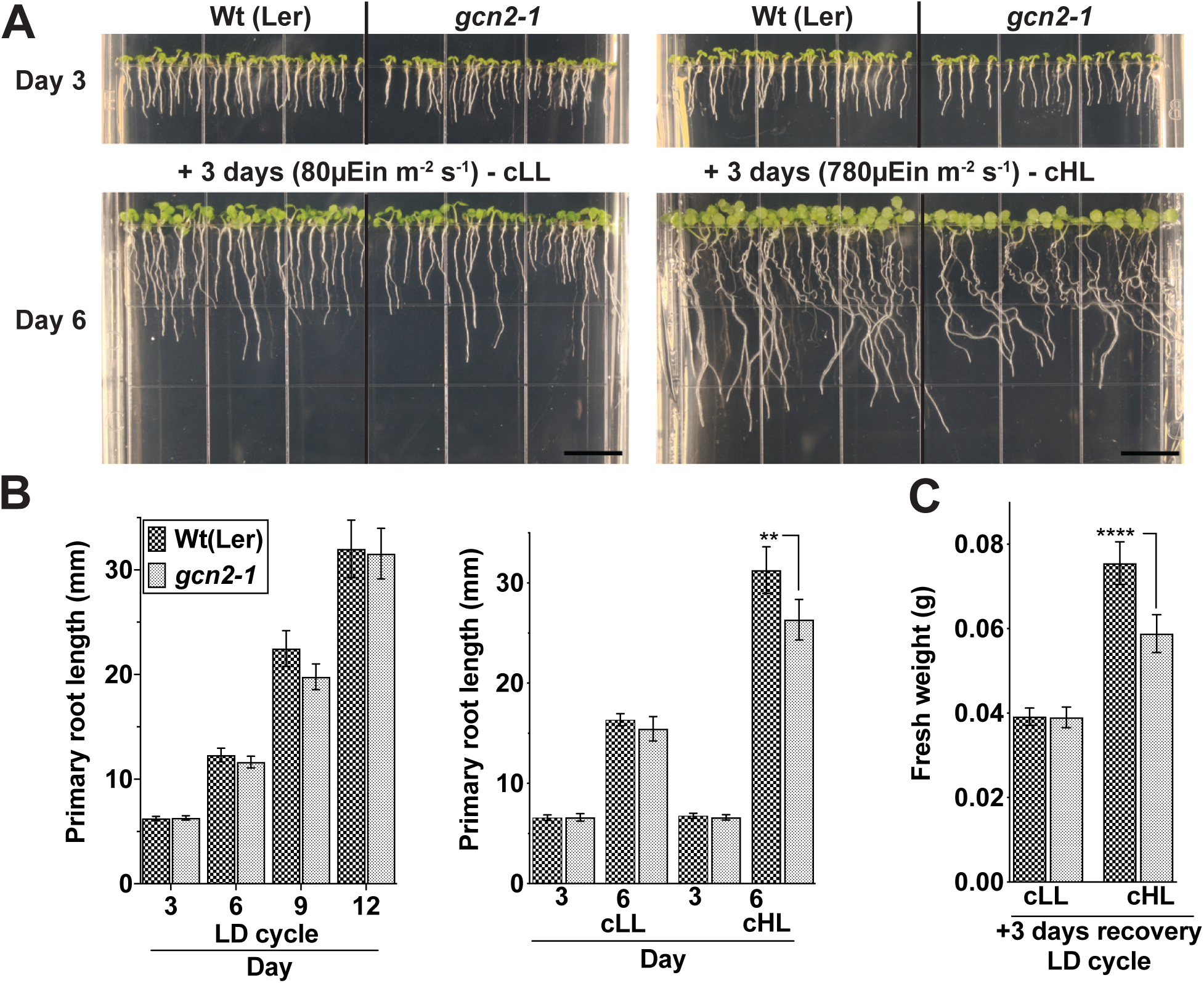
Loss of *GCN2* renders increased sensitivity towards high light stress. **(A)** Top panel, 3-days-old wild-type Landsberg (Wt (Ler)) and *gcn2-1* mutant (*gcn2-1*) seedlings grown under long day period on plant media supplemented with 0.1% sucrose. Bottom panel, the same seedlings after 3 days of additional continuous light at 80 µEin m^-2^s^-1^ (cLL, left panel), or continuous high light at 780 µEin m^-2^s^-1^ (cHL, right panel). Scale bars are 10mm. **(B)** Left - Primary root length of Wt and *gcn2-1* mutants grown under 16 hr light and 8 hr dark (LD) cycle, Right - Root length of Wt and *gcn2-1* as shown in panel (A). **(C)** Seedling fresh weights of Wt and *gcn2-1* mutants from panel (B) after an additional 3 days of recovery in a long day period after the cLL or cHL treatment. Error bars represent standard deviation of four biological replicates with n>12 (B-left) or n>150 (B-right, C) per experiment (Welch’s t-test **P-value <0.005; **** P-value <0.0001).

**Figure 7.**
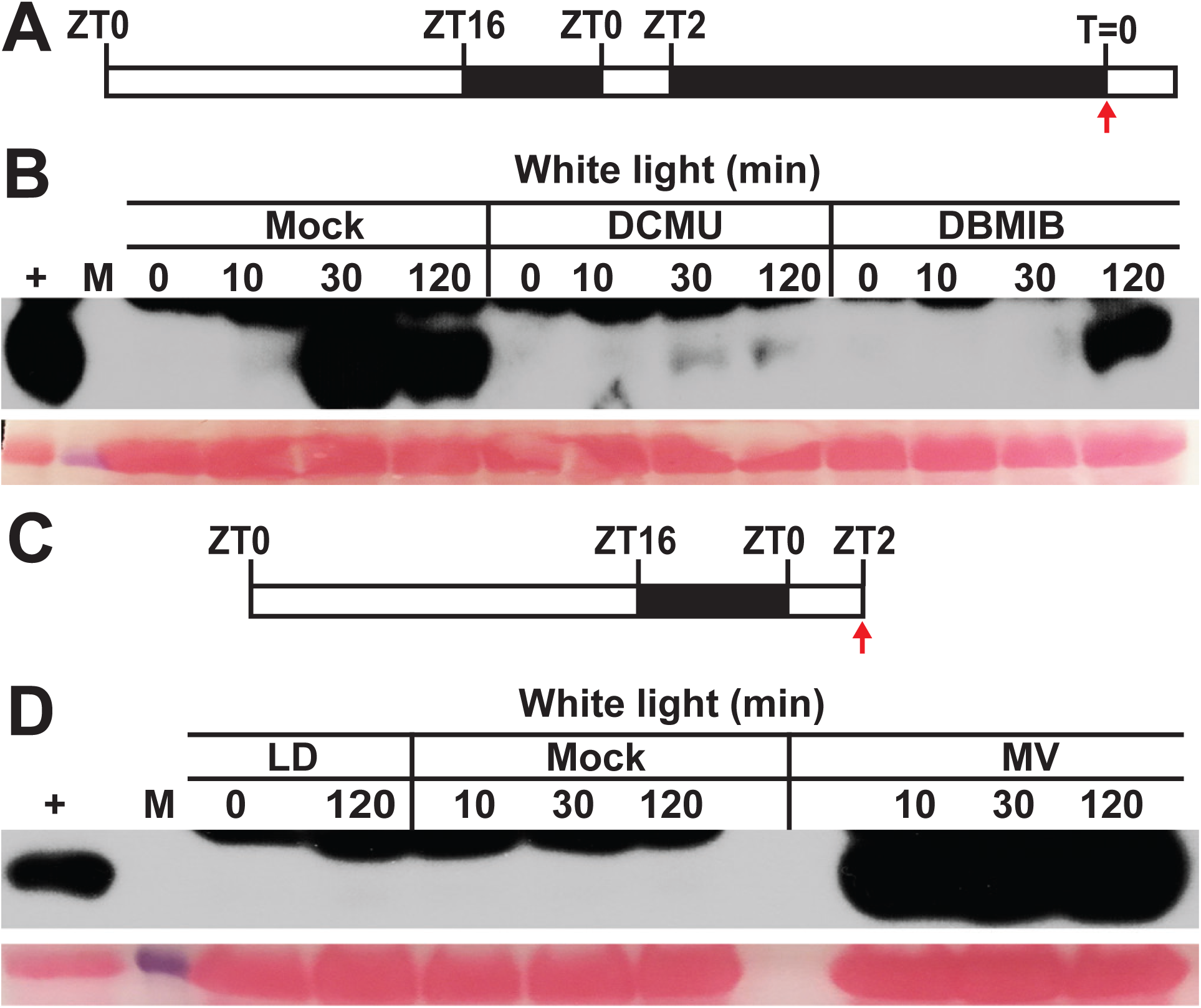
Photosynthetic inhibitors modulate eIF2α phosphorylation in the light. **(A)** Schematic of the light regimen consisting of a long day period followed by 24 hr dark acclimation and the start time of light treatment (80 µEin m^-2^s^-1^) and sampling (T=0). **(B)** Time course of eIF2α phosphorylation in 14-days-old wild-type Landsberg seedlings after lights-on. Thirty minutes prior to light exposure, seedlings were sprayed with either DMSO (Mock), or 8µM DCMU, or 16µM DBMIB. **(C)** Schematic of the light regimen for panel (D). **(D)** eIF2α phosphorylation starting at ZT2 (T=0) either left untreated (LD) or treated with water up to 120 minutes (Mock) or treated with 20µM methyl viologen (MV). For details see legend to Fig.1.

What are the defects of *gcn2* at the level of cellular physiology? The *gcn2-1* mutants had no dramatic differences in photosystem II efficiency during exposure to continuous high light except for a slightly reduced quantum yield overall (**Supplemental Figure 8**). Moreover, in our hands, the *gcn2-1* mutants accumulated as much ROS as wild type under light conditions when GCN2 kinase is active (dark-to-light shift) (**Supplemental Figure 9)**. This result stands in contrast with previously published data, where *gcn2-1* plants accumulated less ROS than wild type in response to treatment with the herbicide, glyphosate (Faus et al., 2015). The *gcn2-1* mutant also did not reveal any difference in survival or overt phenotypes when light-grown seedlings were exposed to a concentration series of H_2_O_2_ (1-1000 mM) or to methyl viologen (data not shown). A drop in polysome loading was observed when WT was treated with H_2_O_2_; however, this drop was not abrogated in the *gcn2-1* mutant (**Supplemental Figure 10**). Therefore, although GCN2 responds to ROS and helps the plant to acclimate to ROS-producing conditions such as high light, it appears to regulate processes other than photosynthesis or H_2_O_2_ levels.

**Figure 8.**
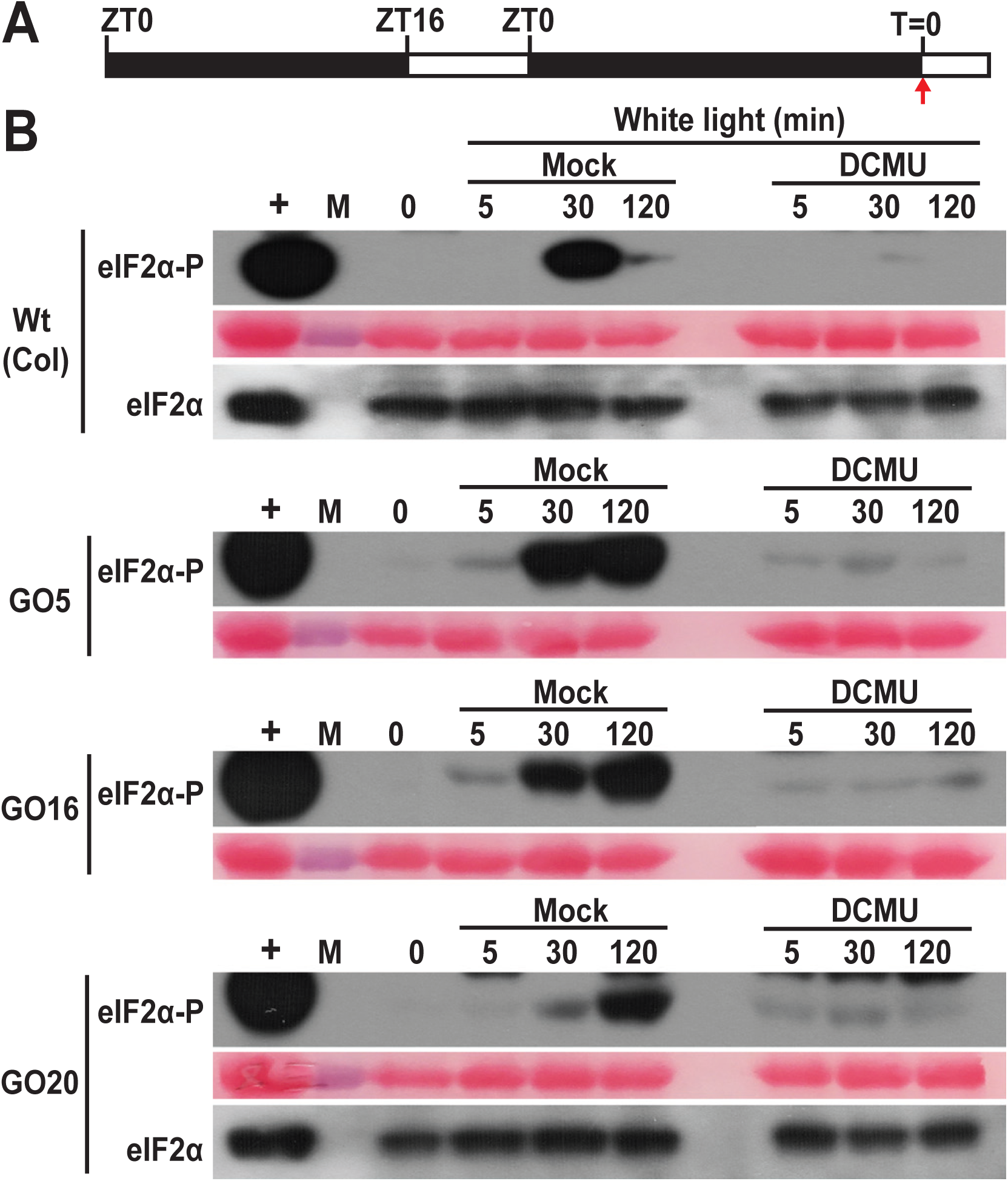
Ectopic glycolate oxidase augments GCN2 activation by photosynthetic ROS. **(A)** Schematic of the light regimen. 30-days-old seedlings grown in a short day cycle were dark adapted and then exposed to light starting at T=0 (red arrow). **(B)** Time course of eIF2α phosphorylation in wild-type Columbia (Wt(Col)) and three independent lines over expressing glycolate oxidase targeted to the chloroplast (GO5, GO16, GO20). Seedlings were sprayed with either DMSO (Mock) or 8µM of DCMU 30 minutes prior to light exposure. For details see legend to Fig.1

**Figure 9.**
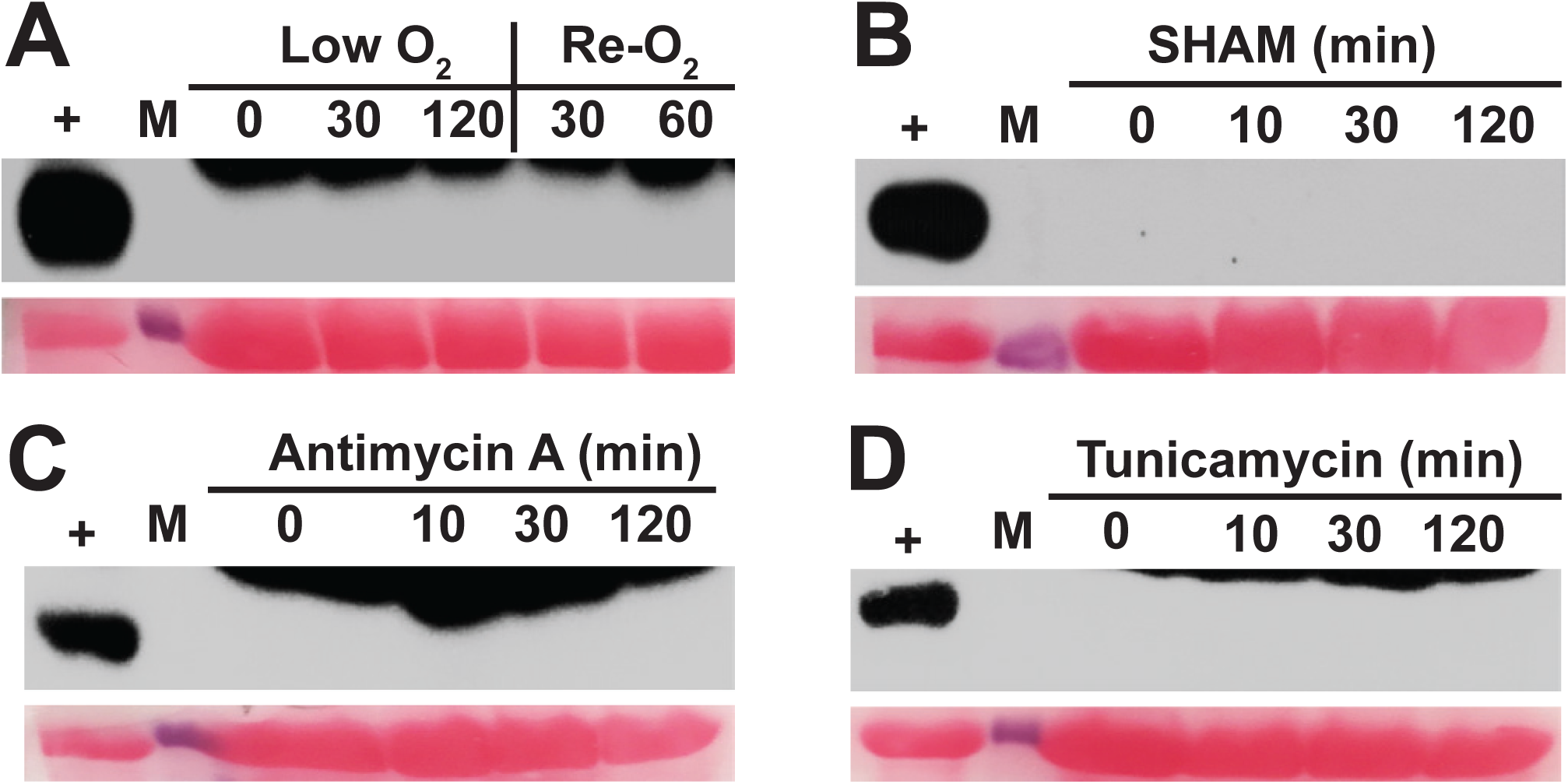
eIF2α phosphorylation is specific to ROS from the chloroplast. 14-days-old wild-type Landsberg seedlings were dark-acclimated for 24 hr and treated as follows to induce ROS stress from the mitochondria or ER. **(A)** Hypoxia (low O_2_) with argon gas followed by re-oxygenation (Re-O_2_) in air. **(B)** Sprayed with 200µM salicylhydroxamic acid (SHAM), an inhibitor of alternative oxidase. **(C)** Sprayed with 50µM antimycin A, a mitochondrial complex III inhibitor. **(D)** Sprayed with 5µg/ml tunicamycin to induce ER stress. For details see legend to Fig 1.

**Figure 10.**
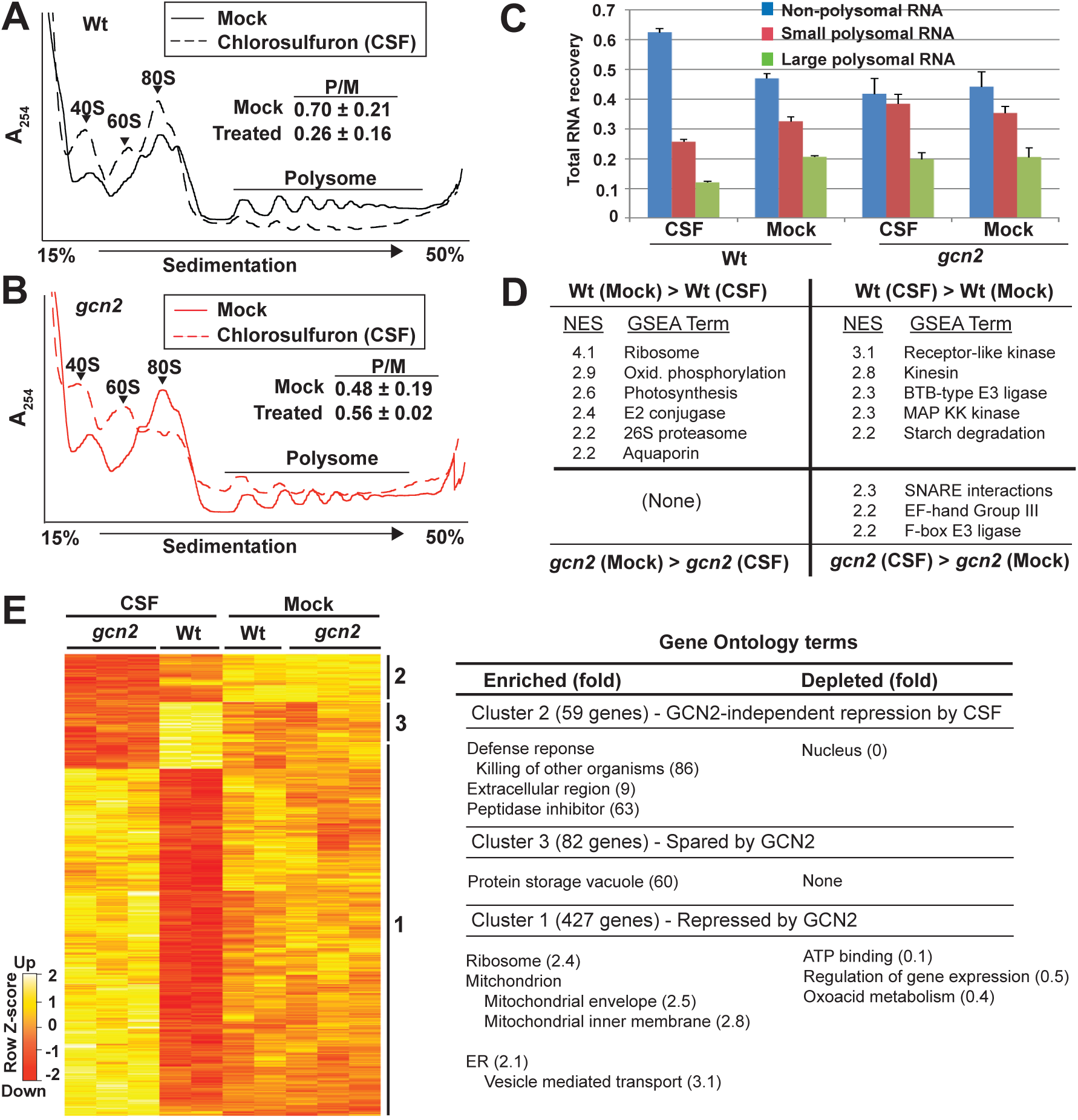
GCN2 regulation of mRNA translation state. Cell extracts from Arabidopsis seedling shoots treated with 0.5µM chlorosulfuron (2 hr) were fractionated over sucrose gradients to separate polysomal from non-polysomal RNAs. **(A, B)** UV absorbance profiles of polysome gradients. Upon activation by CSF, GCN2 kinase inhibits polysome loading in the wild type but not in *gcn2* mutants. The ratio of polysomes (P) to monosomes (M) is indicated with standard error from 3 replicates. **(C)** Gradient fractions were pooled into non-polysomal (NP), small (SP) and large polysomal (LP) RNA pools respectively. The histogram shows the average RNA recovered with standard error. **(D)** Gene set enrichment analysis of mRNA translation states. The differential translations states of 13,551 mRNAs were rank-ordered for each of two pairwise comparisons (Wt±CSF, *gcn2*±CSF). Next, 257 gene sets harboring between 15 and 500 members were examined for a biased distribution along the rank-ordered mRNAs. Gene sets with a bias passing a family-wise error rate < 0.05 are listed with their normalized enrichment score (NES) where 0 equals no enrichment. **(E)** The NP, SP, and LP RNA samples were processed for microarray gene expression profiling. Limma with FDR correction (p < 0.05) identified 568 differentially translated genes. Translation state values are displayed as a Z-score. Three major clusters were identified and analyzed for functional enrichment or depletion using topGO (for details see Supplemental Table 1).

### GCN2 is activated by ROS from the chloroplast but not other organelles

We hypothesized that the activation of GCN2 by light involves ROS produced by photosynthesis in the chloroplast. Strikingly, DCMU almost completely suppressed GCN2 kinase activity under light conditions, while DBMIB appeared to be slightly less effective (**Fig. 7A, B)**. These results indicate that excess light activates GCN2 through photosynthetic electron transfer, and the activation signal likely originates from the PQH_2_ pool. The reason that DBMIB also inhibited GCN2, even though DBMIB should push PQ towards the ROS-producing PQH_2_ form, may lie in the overall reduction in photosynthetic electron flow and hence ROS production when DBMIB is present. In keeping with the hypothesis that photosynthetic ROS activates GCN2, norflurazon, a herbicide that inhibits carotenoid biosynthesis (Sandmann, 1994), rapidly activated GCN2, which was kept in check by pretreatment of seedlings with DCMU or DBMIB (**Supplemental Figure 11**). Similarly, the herbicide methyl viologen (MV, paraquat), which leads to rapid superoxide-mediated hyper-accumulation of H_2_O_2_ in chloroplasts (Fujii et al., 1990), triggered intense GCN2 activity within 10 min (**Fig. 7C, D**). As expected, methyl viologen stimulated (**Fig. 4E**) and DCMU suppressed ROS production by light *in situ* (**Fig. 2D**). MV did not activate GCN2 under dark conditions (**Supplemental Figure 12**). However, MV significantly lowered the photosystem II quantum yield of the *gcn2-1* mutant compared to wild-type plants (**Supplemental Figure 13**), suggesting a deficiency in recovery from ROS stress in the *gcn2-1* mutant that may potentially underlie the root growth defect evident under high light.

**Figure 11.**
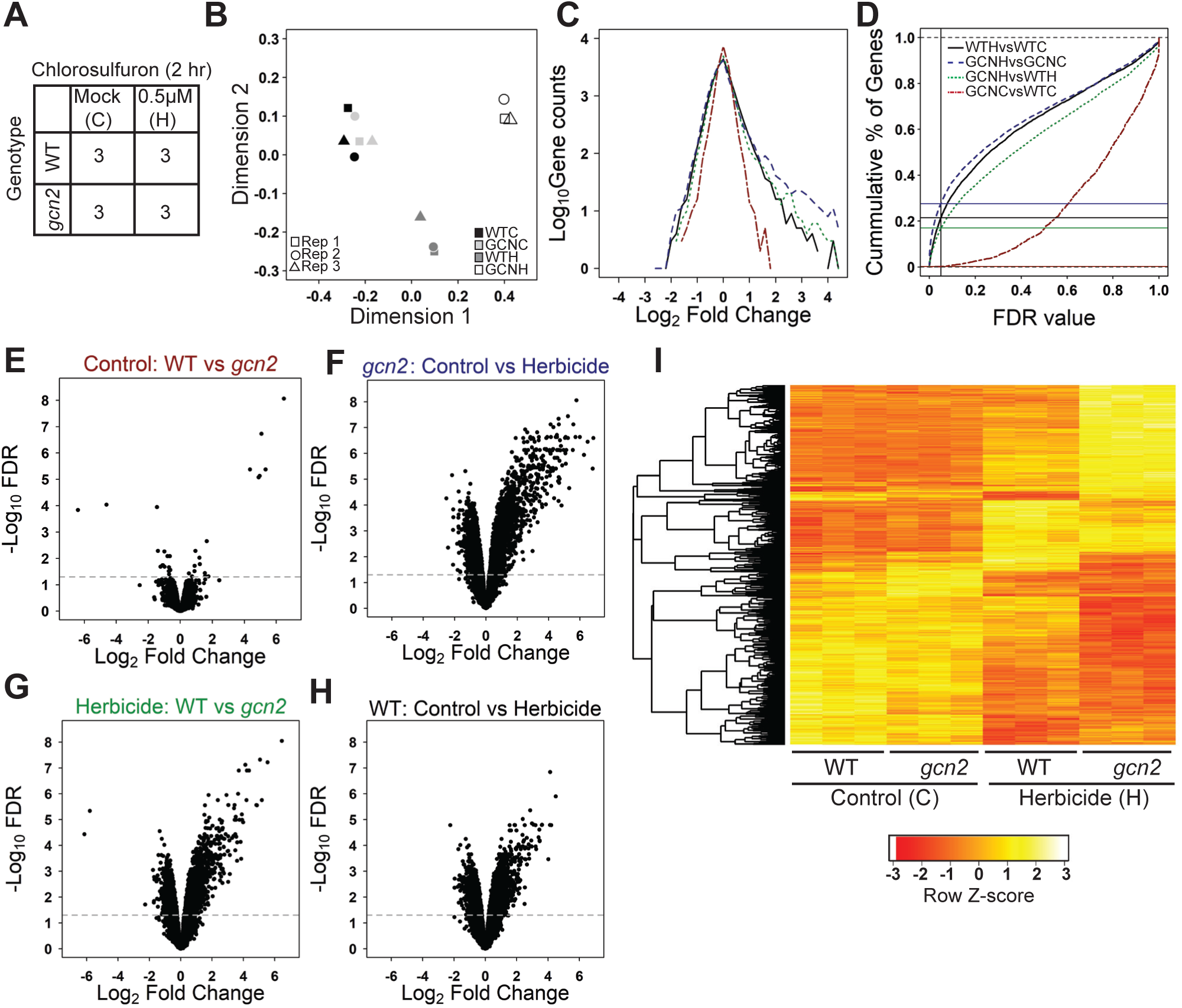
Transcriptome analysis of *gcn2* mutant seedlings under chlorosulfuron herbicide. **(A)** Schematic of the experiment. The ‘3’ indicates three biological replicates. **(B)** Gene expression data are displayed by multidimensional scaling. Replicates are identified by a square, circle and triangle. C and H stand for control and herbicide-treated. The plot was produced using limma’s plot MDS function on all the data. **(C)** Line histogram of the number of differentially expressed genes for the four pairwise comparisons described in (D). See panel (C) for color codes. **(D)** Line graphs of the number of genes deemed differentially expressed as a function of the false discovery rate for the four pairwise comparisons. An FDR of 0.05 is marked by the vertical line. The data were corrected for false discovery by multiple comparisons using the Benjamini-Hochberg method. Red, control-treated *gcn2* versus control-treated WT. Green, herbicide-treated *gcn2* versus WT. Black, WT control versus WT herbicide; Blue, *gcn2* control versus *gcn2* herbicide. The horizontal lines highlight where each curve intercepts FDR=0.05. **(E-H)** Scatter plots (‘volcano plots’) of the fold-changes in mRNA level versus the false-discovery rate (FDR) for the four pairwise comparisons in (D). The horizontal line marks FDR=0.05. **(I)** Cluster analysis of the four experimental transcriptomes. Included are 4632 genes (probe sets) that showed significant differences in mRNA level in both a one-way ANOVA and any pairwise comparison (limma FDR<0.05). The clustering tree is indicated on the left.

**Figure 12.**
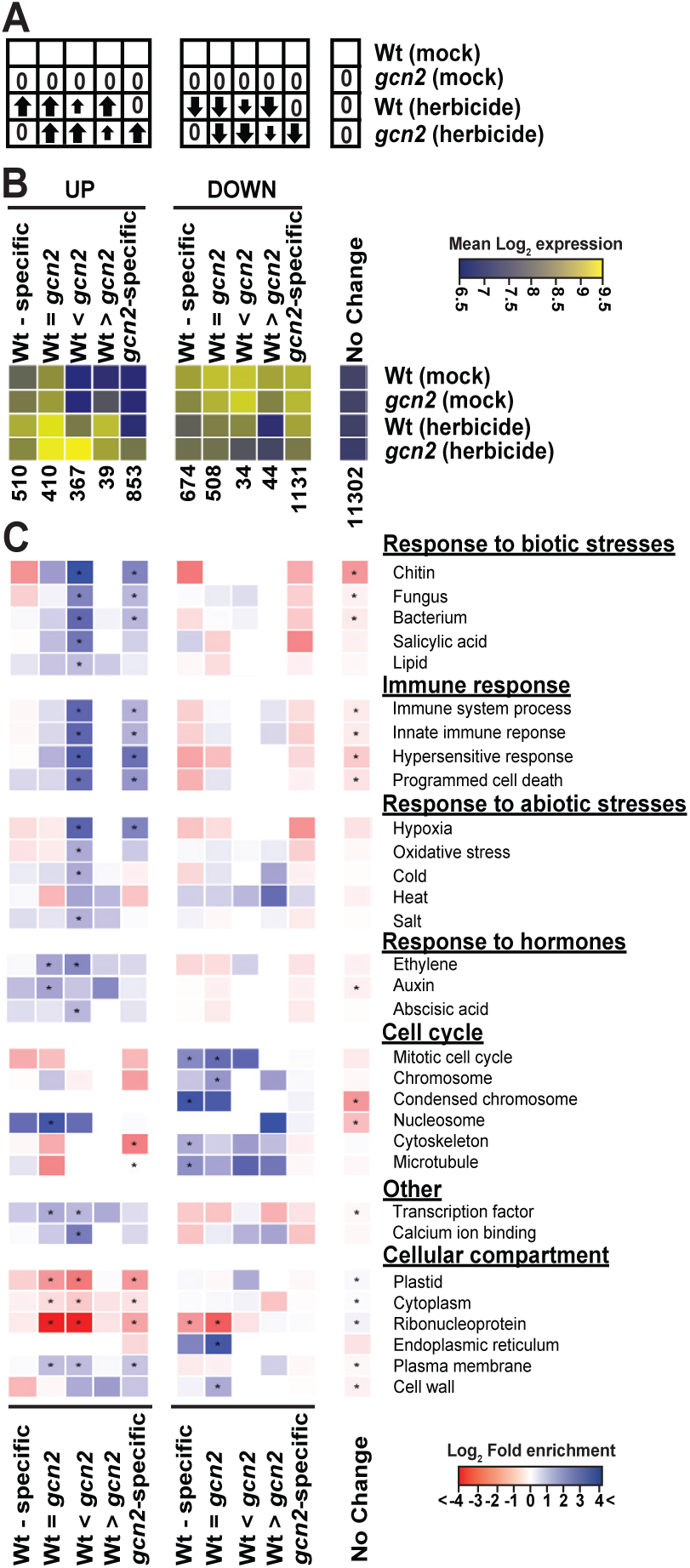
Gene ontology analysis of the GCN2-responsive transcriptome. **(A)** Cartoon illustrating the filter being applied under each of eleven bins. An arrow indicates whether transcript levels are up or down or unchanged (0) compared to mock-treated wild-type Landsberg (Wt) as a reference. The small arrow represents a lesser response. **(B)** Heatmap of the average mRNA levels in each of the eleven bins in (A). The color scale shows the mean log2-transformed hybridization signals per bin on the Affymetrix gcRMA scale. The number of genes in each bin is indicated. Note that genes unresponsive to herbicide typically have lower mRNA levels than herbicide-responsive genes. **(C)** All eleven bins were analyzed by gene ontology with topGO with correction for multiple testing. The color-coded heatmap displays the fold-enrichment of a given gene ontology term in each bin over what is expected by chance. Blue hues denote enriched terms and red hues are depleted terms. Cells where the enrichment/depletion passes FDR <0.05 are labeled with an *.

Plants that overexpress a chloroplast-targeted glycolate oxidase (GO), a peroxisomal enzyme, provoke plastid-specific H_2_O_2_ responses in Arabidopsis (Fahnenstich et al., 2008). GO overexpressing seedlings of three independent lines were dark-adapted and then exposed to light. These seedlings showed slightly accelerated and sustained GCN2 activation compared to the corresponding wild-type Columbia (**Fig. 8A, B**), which could be suppressed with DCMU (**Fig. 8B**), consistent with the elevated ROS accumulation in the GO overexpressing seedlings (Fahnenstich et al., 2008). Taken together, these observations support that light stress leads to rapid production of H_2_O_2_ in chloroplasts, and this may serve as the activation signal for GCN2.

Next, we asked whether GCN2 can be activated by ROS generated from organelles other than the chloroplast. We treated hypoxic seedlings with oxygen, and we treated normal seedlings with the mitochondrial electron transport inhibitor antimycin-A (Maxwell et al., 1999) and the alternative oxidase inhibitor salicylhydroxamic acid (SHAM) (Stonebloom et al., 2012), both of which trigger mitochondrial ROS in the dark (**Supplemental Figure 14**). Neither of these treatments activated GCN2. Likewise, tunicamycin, an inhibitor of protein folding in the ER that can trigger ROS accumulation (Cao and Kaufman, 2014; Ozgur et al., 2014) (**Supplemental Figure 14**), was also inactive towards GCN2 under dark (**Fig. 9A-D**). Taken together, by manipulating the redox status of the chloroplast PQ pool or other photosynthetic components, we were able to control eIF2α-P levels.

### How does GCN2 regulate mRNA ribosome loading?

To investigate the consequence of GCN2 kinase activation at both the translational and transcriptional levels, we activated GCN2 in light-grown seedlings using the herbicide chlorosulfuron. Cell extracts from wild-type and *gcn2* seedling shoots were fractionated on sucrose gradients to separate polysomal from non-polysomal RNAs. As expected, (Lageix et al., 2008; Zhang et al., 2008), GCN2 globally inhibited polysome loading in response to the brief (2 hr) herbicide treatment (**Fig. 10A-C**). Following microarray hybridizations of non-polysomal, small polysomal and large polysomal mRNAs, we calculated an *ad-hoc* ‘translation state’ (TL) for each mRNA (Missra et al., 2015) in each of the four treatments (i.e., ± GCN2 and ± herbicide). We identified differentially translated mRNAs (**Fig. 10E** and **Supplemental Table 1**). Wild type and *gcn2-1* plants had similar translation states under control conditions (**Fig. 10E**), in keeping with the lack of kinase activity and the weak phenotypic differences. After herbicide treatment, three major response types were evident. Cluster 1 was the largest and was preferentially repressed by active GCN2 kinase. Proteins encoded here were enriched for three cellular locales, ER and vesicle trafficking, ribosomes, and mitochondrial membrane complexes, in particular proton-coupled ATP synthesis. Interestingly, general ATP binding and phosphorylation (e.g. kinases) were strongly depleted from this cluster, indicating that many remained preferentially ribosome-loaded. Cluster 2 was translationally repressed to a similar degree in wild type and *gcn2-1*. The proteins encoded in this herbicide-repressed but *bona fide* GCN2-independent cluster were enriched for extracellular proteins and defense responses and were strikingly depleted for nuclear functions. Cluster 3 encodes mRNAs that were spared from the translational repression by active GCN2 and was enriched for only few functional terms (**Fig. 10E**), but not surprisingly included the mRNA for GCN2 kinase itself. For a more sensitive functional analysis that surveys the entire translatome rather than just the few mRNAs passing a stringent false discovery rate, we turned to gene set enrichment analysis (GSEA, **Fig. 10D**). GSEA of the herbicide response strongly confirmed some of the earlier trends (repressed: ribosome, mitochondrial oxidative phosphorylation; spared from repression: kinases), but also revealed new trends, especially in the protein turnover domain, as well as translational repression of photosynthesis (thylakoid) functions. GSEA also revealed functional biases in the milder, GCN2-independent translational control by herbicide. Thus, generally, core metabolic machinery was especially strongly repressed by GCN2, while regulators such as kinases and E3 ligases were relatively protected from translational repression.

To interpret these data properly we note that wild-type plants experience a global reduction in polysome loading under our treatment conditions; this global reduction is masked here because the array hybridization was performed with equal amounts of total RNA per sample. We then reanalyzed the wild-type translatome by unmasking the global translational repression by herbicide in the wild type (**Supplemental Table 2**). This analysis further confirmed the gist of the gene ontology from clustering and GSEA, yet also uncovered additional details that link the molecular physiology of the GCN2 pathway with functional domains at the level of translational control. For example, relatively protected from translational repression were hydrogen peroxide metabolism (trend), ethylene metabolism, defense responses, and aspects of amino acid homeostasis.

### GCN2 remodels the transcriptome, implicating GCN2 in plant responses to stress

Although GCN2’s primary known role is as a translational regulator, one would expect that alterations in translation will result in subsequent alterations in mRNA transcript levels, which in turn may shed light on the role of GCN2 at the physiological scale (**Fig. 11A** and **Supplemental Dataset 2**). The four sets of triplicate transcriptome data separated well in a multidimensional scaling plot (**Fig. 11B**). In the absence of herbicide, wild type and *gcn2* mutants had nearly identical transcript levels (**Fig. 11C, D, E**), in keeping with the nearly indistinguishable whole-plant phenotypes. The response to herbicide was dramatic and was biased towards upregulation in both WT and *gcn2* (**Fig. 11C, F, H**), as expected given that the herbicide treatment was short compared to the average lifetime of mRNAs. Many individual genes responded more strongly in the *gcn2* mutant (blue traces) than in WT (black traces) (**Fig. 11C-D, F-I**). Hence, the *gcn2* mutant is hypersensitive to the herbicide stimulus.

Although unsupervised clustering of the transcriptome response revealed clearly delineated response types (**Fig. 11I**), we decided to bin genes according to 11 predefined filters (**Fig. 12**) for a finer-grained and biologically motivated classification. The herbicide response was strongly affected by the *GCN2* genotype, ranging from genes that only responded in wild type and not in *gcn2* (WT-specific) all the way to genes that only responded in *gcn2* but not in wild type (*gcn2-* specific, **Fig. 12A, B**). We were most interested in the gene ontology of those mRNAs that respond to herbicide in WT and whose response was exaggerated in *gcn2* (columns 3 and 8 in **Fig. 12B, C**), indicating that wild-type GCN2 tempers their response to herbicide. Among these, the upregulated genes were biased towards “response to stimulus” (201/367), especially biotic defense and immune responses, as well as many abiotic stresses including oxidative stress. This group includes genes for ROS signaling, for example, the famous *RbohD* isoform of the *NADPH oxidase* that is responsible for the respiratory burst. The strongest enrichment was seen for “response to chitin”, a gene ontology term that contains numerous transcription factors. Thus, herbicide treatment triggered a defense response, which was attenuated by active GCN2 kinase. Another bin comprised genes whose innate response to herbicide was visible in the *gcn2* mutant but completely suppressed by GCN2 (column 5, *gcn2*-specific). This bin was enriched for similar terms as the previous one and thus appears to be part of the same regulatory mechanism.

Several functional groups were strongly depleted among the herbicide-responsive genes, namely the plastid-bound transcriptome as well as ribonucleoprotein complexes, i.e. translation and ribosome biogenesis. Given that chlorosulfuron inhibits amino acid synthesis in the chloroplast and thus translation of chloroplast proteins and potentially retrograde signals from the chloroplast to the nucleus, it is surprising how little the plastid-bound transcriptome has responded.

### GCN2 mediates chitin priming

Exposure to chitin primes an innate immune response that protects plants against the bacterial pathogen *Pseudomonas syringae* DC3000 (Zipfel and Robatzek, 2010). Given that ‘response to chitin’ and related immune responses were the most prominent functional categories among the mRNAs that are upregulated in the *gcn2-1* mutant, we next examined whether *gcn2-1* mutants respond differently to chitin. Interestingly, while defense priming by chitin was observed in wild-type Landsberg, this pattern triggered immunity (PTI) response was significantly reduced in the *gcn2-1* mutant (**Supplemental Figure 16**), indicating that GCN2 kinase plays a role in PTI to some degree and the activation of innate immune signaling. On one hand, the loss of priming by chitin in the *gcn2-1* mutant is somewhat surprising because transcript levels in the ‘response to chitin’ category were elevated rather than reduced in *gcn2-1* treated with herbicide (**Fig. 12C**). However, this observation may indicate that the *gcn2-1* mutant response to herbicide triggers a complex series of events that ultimately link the GCN2 kinase activity with cellular signaling processes associated with stress, including biotic stress responses.

## DISCUSSION

Extensive studies in animals and yeast demonstrate that the GCN2 kinase mediates global translational repression in response to environmental signals by phosphorylating its primary substrate, eIF2α (Dever et al., 1992). There, the GCN2 kinase is activated by uncharged tRNA (Wek et al., 1989; Wek et al., 1995; Dong et al., 2000; Anda et al., 2017). In plants, the GCN2 pathway appears to be substantially conserved; specifically, GCN2 is encoded by a single gene in Arabidopsis (Zhang et al., 2003). GCN2 kinase can be activated by uncharged tRNA *in vitro* (Li et al., 2013). *In planta*, the kinase is activated by inhibitors of amino acid biosynthesis such as the herbicides chlorosulfuron, glyphosate and glufosinate (Lageix et al., 2008; Zhang et al., 2008), and it is activated in a mutant with a defect in cysteine biosynthesis (Dong et al., 2017). GCN2 kinase phosphorylates eIF2α (Lageix et al., 2008; Zhang et al., 2008), and the inhibitors of amino acid biosynthesis that activate the GCN2 kinase indeed reduce overall ribosome loading of mRNAs (Lageix et al., 2008).

### GCN2 activation by light and ROS

The true nature of the biochemical signal that activates GCN2 *in planta* remains unclear. Herbicides are synthetic activators of GCN2. The GCN2 kinase is activated by numerous other agents, including ultraviolet light, wounding, the ethylene precursor 1-ACC, and the endogenous defense signals salicylic acid and methyl-jasmonate (Lageix et al., 2008), but it is unclear whether these signals are the authentic activators for GCN2 in nature. Here we describe that the GCN2 kinase is also activated by light in a dosage dependent fashion. The activation is seen most clearly in 24 hr-dark adapted plants, where GCN2 activity becomes undetectable within 6 hr of dark acclimation and then increases rapidly upon exposure to excess light stress. Some activation of GCN2 is also evident in plants grown under a regular long-day light-dark cycle. Significantly, even the traditional activation of GCN2 by herbicides is strictly light dependent. These observations suggest that, beside uncharged tRNA as a ligand for the kinase, a second, light-dependent signal is required.

It is unclear which biochemical sequence of events causes excess light to activate GCN2. The light-dark pattern of GCN2 activation coincides with the redox rhythm. Our experiments suggest that ROS, which are produced in the chloroplast during photosynthesis, may be a key intermediate, although we do not rule out that uncharged tRNA may also be required. First, GCN2 is activated rapidly by reactive oxygen species, by application of hydrogen peroxide ectopically, and even more rapidly by methyl viologen, which triggers superoxide and hydrogen peroxide production from photosystem I (Fuerst and Norman, 1991). Internal triggers of plastidic ROS, i.e. plastidic expression of glycolate oxidase, application of norflurazon, and exposure of *flu1* mutants to excess light, also stimulate GCN2 activity. In keeping with a role for ROS, the activation of GCN2 by excess light can be suppressed with ascorbate. It can also be repressed by manipulating the redox status of the photosynthetic apparatus with electron transport inhibitors, suggesting that a plastidic ROS is a source of the signal that activates GCN2. Chemical or other treatments that stimulate ROS production in the mitochondria (Stonebloom et al., 2012; Paradiso et al., 2016; Cui et al., 2018) did not activate GCN2 at the time periods tested, nor did tunicamycin, a trigger of ER stress well known to cause eIF2α-P in animals (Leitman et al., 2014).

It has long been known that ROS produced in the chloroplast can regulate translation in the cytosol (Khandal et al., 2009), with methyl-jasmonate as a potential intermediate. Our work suggests that this pathway may operate in part via the GCN2 kinase. That methyl-jasmonate can activate GCN2 (Lageix et al., 2008) supports this hypothesis. However, in our hands, ROS also repressed translation in a GCN2-independent manner. These observations suggest that H_2_O_2_ may regulate cytosolic translation by multiple pathways. A similar situation exists in fission yeast where H_2_O_2_ stimulates the GCN2 kinase and represses translation, but the effect on translation is at least partially GCN2-independent (e.g. Knutsen et al., 2015). These comparisons suggest that ROS-mediated GCN2 kinase activation in Arabidopsis is evidence of a pan-eukaryotic phenomenon. The mechanism whereby ROS activates the GCN2 kinase remains unknown. Given the speed at which GCN2 is activated, the most likely candidates are a direct biochemical activation of the kinase by a redox-active compound or by an intermediate that is released from the chloroplast during ROS stress. This very question also remains unresolved in every other organism, fission yeast, budding yeast, mammals, where ROS-activation of eIF2α kinases has been observed over the past 15 years (Shenton et al., 2006; Anda et al., 2017) and remains to be explored in more detail.

### The effect of GCN2 on the translatome and transcriptome

Given that the overt growth defects in the *gcn2* mutant were mild, we analyzed its transcriptome to identify the consequences of GCN2 activation. We decided to stimulate GCN2 kinase activity with herbicide rather than ROS because, first, ROS have numerous biochemical targets and trigger multiple signaling pathways, and second because ROS accumulate in a non-uniform pattern across tissues. In contrast, chlorosulfuron herbicide specifically inhibits branched-chain amino acid synthesis. In addition, chlorosulfuron suppresses bulk ribosome loading in a GCN2-dependent manner (Lageix et al., 2008), whereas hydrogen peroxide did not, under our conditions. Similar to data by (Faus et al., 2015), in the absence of herbicide the transcriptomes of WT and *gcn2* are essentially the same. Once GCN2 kinase is activated by herbicide, two roles can be distinguished. The first of these is “GCN2 as attenuator”; among all the herbicide-inducible mRNAs, GCN2 attenuates more than half, either partially or completely. Many of these mRNAs are functionally associated with responses to biotic or abiotic stresses (**Fig. 12C** column 3 and 5). The remaining herbicide-inducible mRNAs are not affected by GCN2. What distinguishes these at the molecular level is currently unknown, but there appears to be a difference because the GCN2-independent mRNAs are significantly enriched for ‘response to auxin’ but less enriched for responses to stress. The second role of GCN2 kinase is “GCN2 as repressor/activator”; after treatment with herbicide, many mRNAs become repressed, often with the help of GCN2 (**Fig. 12C** column 6 and 9), and some of these are cell cycle related. Taken together it is noteworthy that a substantial portion of the transcriptomic response to herbicide appears to be mediated by the GCN2 kinase.

The *gcn2* mutant plants have an exaggerated defense response. Can this be explained by the loss of translational control in the *gcn2* mutant? In detail, in the wild type, the herbicide treatment causes changes in transcript levels, but the herbicide also represses translation via GCN2. In *gcn2*, those changes are exaggerated, possibly because translation of specific regulatory mRNAs remains high. Therefore, we also analyzed the translatome of GCN2 under the same herbicide treatment. GCN2 changed the translation state of hundreds of mRNAs. Thus, it stands to reason that the transcriptional role of GCN2 as an attenuator of defense responses is mediated by its translational repressor function. How? It is not possible to attribute the effect to a single mRNA. However, we know that about twenty of the mRNAs that are hyper-translated in *gcn2* code for likely transcriptional regulators (**Supplemental Dataset 1**, Cluster 1). In animals and budding yeast, GCN2’s translational effects on mRNAs such as ATF4 and GCN4 and the resulting transcriptional changes are known as the ‘integrated stress response’ and the ‘general amino acid control’, respectively (Wek, 2018). For comparison, in fission yeast a non-homologous but functionally analogous mRNA for a GATA transcription factor was only recently identified (Duncan et al., 2018). Arabidopsis appears to make use of GCN2 kinase in a different way, but eventually tying a stress signal, in our case herbicide, to a transcriptional response. In budding yeast and animals, activation of GCN2 will boost the ribosome loading of certain bZip-type transcription factors that harbor uORFs in their 5’ untranslated region. Although bZips and several other classes of transcription factors harbor uORFs in Arabidopsis (von Arnim et al., 2014), we did not observe that these mRNAs are in any way poised for elevated translation once GCN2 is activated. This result may indicate that GCN2 and eIF2α-P affect translation differently in plants than in yeast and animals.

### The physiological role of GCN2 kinase

One conundrum surrounding plant GCN2 is that *gcn2* mutants have rather mild phenotypes under favorable lab conditions (e.g. Liu et al., 2015) and a near-normal transcriptome. Although the GCN2 kinase can be activated by numerous treatments, very few of these conditions cause a maladaptive phenotype in *gcn2* mutants, lending urgency to the question what signals activate GCN2 in the wild. We show here that the GCN2 kinase is not only activated by excess light but *gcn2* mutants also have a mild growth deficiency under prolonged continuous or high light, depending on the ecotype. Seedlings that had been transferred from dark to light also trended towards a GCN2-mediated reduction in polysome loading (**Supplemental Figure 15**). These data support the notion that adaptation to high light is part of the functional portfolio of the GCN2 kinase.

Our data add to the evidence that GCN2 is part of the biotic defense response. First, GCN2 is activated by signals that arise during regular defense responses such as salicylic acid, jasmonic acid, ethylene (Lageix et al., 2008) and reactive oxygen (our data). GCN2 is also activated in some but not all settings of bacterial infections (Izquierdo et al., 2018; Liu et al., 2019). Second, the transcriptome of *gcn2* overexpresses mRNAs from defense related gene ontology terms such as “response to chitin” and the oxidative burst. Thus, the presence of GCN2 can attenuate the defense response at the transcriptional level, akin to what is observed during the preinvasive stages of bacterial infection (Liu et al., 2019). Third, GCN2 also preferentially affects the translation of mRNAs in the secretory system, and the secretory system is responsible for secretion of many defense related proteins such as peptidases and chitinases. Finally, *gcn2* mutant plants had a defect in the priming of their bacterial defense response by chitin (our data), but were more resistant to unprimed infection by *Pseudomonas syringae* (Liu et al., 2019). In summary, findings presented here bolster the view that GCN2 is an integral part of the defense response against pathogens. Future studies aimed at identifying changes in the translatome, specifically in a time resolved manner during bacterial infection may shed more light on the plant defense strategies utilizing the conserved GCN2-eIF2α paradigm.

## MATERIALS AND METHODS

### Plant materials and growth conditions

Seeds of *Arabidopsis thaliana* ecotype Landsberg (Ler-0), Columbia (Col-0), and homozygous *gcn2-1* mutants of the GT8359 Genetrap line (Zhang et al., 2008) were sterilized and stratified at 4°C for 2 days. Seeds were germinated on ½-strength Murashige-Skoog (MS) salt plant media (MP Biomedicals, cat # 2633024) with 0.65% Phytoagar (Bioworld, cat # 40100072-2) under a long-day cycle of 16 hr light (80±10 µEin m^-2^s^-1^)/8 hr dark at 22 °C and 50% humidity. Unless stated otherwise, seedlings were grown on media without sucrose. The *flu1-1* mutant is in Landsberg (Meskauskiene et al., 2001) and the glycolate oxidase overexpressing plants are in the Columbia ecotype (Fahnenstich et al., 2008).

The *GCN2* T-DNA insertion mutant (SALK_032196) and (SALK_129334.2) (Alonso et al., 2003), henceforth called *gcn2-2 and gcn2-3* respectively, were obtained from the Arabidopsis Biological Resource Center (ABRC) and characterized by PCR genotyping. Genomic DNA was extracted from 2-week-old *gcn2* mutant seedlings using the GeneJET Plant Genomic DNA purification kit (ThermoFisher, cat# K0791) as per manufacturer’s instructions, and PCR was performed with gene specific *GCN2* primers (GCN2-F1: 5′-AATTCGCCAAATTGTGGAAG-3′, GCN2-R1: 5′-ATAAGCAAATGACAGGTCCG-3′) for *gcn2-2* and (GCN2-F2: 5′-TAAGTTCCCCTGTGTCCCAC-3′, GCN2-R2: 5′-ACTTGGAGACATCAAACGCC-3′) for *gcn2-3*, and a left border T-DNA primer (LBa1: 5′-TGGTTCACGTAGTGGGCCATCG-3′). The predicted T-DNA insertion site was verified by DNA sequencing. Like *gcn2-1, gcn2-2* and *gcn2-3* homozygotes were unable to phosphorylate eIF2α.

### Stress treatments

For stress treatments performed in the dark, 2-week-old seedlings were dark acclimated for 24 hr starting at Zeitgeber time 2 (ZT2). Samples were collected under green safe light. For treatment with H_2_O_2_ (Sigma, cat# H1009), antimycin A (Alfa Aesar, cat# AAJ63522LB0), SHAM (Thermo-Fisher, cat# AC132620050), tunicamycin (Sigma, cat# T7765) and herbicide, dark-acclimated seedlings were sprayed 6-8 times with the respective reagents from a distance of 4 inches. Seedlings were mock treated with either water only or with 0.1% DMSO, as appropriate. For heat and cold stress, plates containing dark acclimated seedlings were shifted to 37 °C or 4 °C respectively in the dark for the desired times. Hypoxia (i.e., low O_2_) stress was administered by using argon gas, and re-oxygenation treatment was performed as previously described (Lokdarshi et al., 2016).

For stress treatments that involved a dark-to-light shift, 2-week-old horizontally grown seedlings were dark acclimated for 24 hr starting at ZT2. A sample for time zero (T0) was collected under green safe light, and for light stress experiments plates were exposed to different light intensities for the desired times. For chemical treatments with DCMU (Thermo-Fisher, cat# D2425), DBMIB (Thermo-Fisher, cat# 271993), chlorosulfuron (Sigma, cat# 34322), glyphosate (Bioworld, cat# 30632003-1), glufosinate ammonium, seedlings were treated (i.e., sprayed) with reagents or mock control (DMSO or water) 30 min prior to the end of the 24 hr dark acclimation. For tests involving antioxidant (i.e., ascorbate), seedlings were grown for 2 weeks on 0.5mM ascorbate supplemented media. Stress treatments with methyl viologen (MV) and norflurazon were performed by spraying 2-week-old seedlings with 20µM MV or 50µM norflurazon respectively.

For pathogen-associated molecular pattern (PAMP)-mediated immunity (PTI) priming, 4-week-old Arabidopsis plants were hand-infiltrated with flg22 (1 µM) or chitin ([GlcNAc]_7_; 1 µM) and incubated at room temperature (∼22 °C) for 24 hr. After 24 hr, PAMP-infiltrated leaves were hand-infiltrated with *Pst* DC3000 strains harbouring the vector pVSP61. For *in planta* bacterial growth enumeration, *Pst* DC3000 strains were inoculated using a needleless syringe at a final concentration of 2 × 10^5^ CFU/ml. Samples were collected at 0 and 96 hr post-inoculation (hpi). All pathogen-inoculation experiments were performed at least three times with three technical replicates per experiment. Statistical analysis of bacterial growth was by one-way analysis of variance using GraphPad Software (Prism).

### Phenotype characterization and high light stress treatment

For phenotypic characterization under high light stress, wild type and *gcn2* mutant seedlings were germinated on ½ strength MS media containing 0.1% sucrose under a long-day cycle of 16 hr light (80±10 µEin m^-2^s^-1^)/8hr dark at 22 °C. Plates with 3 or 4-days-old vertically grown seedlings were transferred to continuous high light (780±10 µEin m^-2^s^-1^) for three consecutive days. For mock treatments, plates were transferred to continuous light (80±10 µEin m^-2^s^-1^) at 22 °C. In order to compensate for any increase in plate temperature as a result of the high intensity fluorescent tube lights (6000 lumens, Super Bright White), chamber temperature was adjusted to 18 °C for the period of high light stress. Additionally, to avoid high light exposure to roots, the vertical plates were covered with black paper and the height of the plates was adjusted to focus light treatment primarily on the shoots. For recovery treatments, both mock and high light exposed plates were transferred to a long day cycle of 16 hr light (80±10 µEin m^-2^s^-1^)/ 8 hr dark at 22 °C for three days before fresh weight measurements.

### Root length, fresh weight measurements and statistical analysis

Photographs of vertically grown seedlings at day 3 post-germination and after stress treatments were taken with a digital camera (Canon). The primary root length was measured using ImageJ (ver. 1.41; http://rsb.info.nih.gov/ij/index.html). For fresh weight measurements, seedlings were weighed on an analytical balance. Statistical tests were performed using GraphPad Prism (ver. 7.0a; GraphPad Software, Inc).

### Protein extraction and immunoblot analysis

Total protein extraction was performed as described previously (Zhang et al., 2008). Briefly, 2-week-old whole seedlings were harvested after the desired treatments and flash frozen in liquid nitrogen. For total protein extraction, seedlings were ground using a plastic pestle in a 1.5 mL tube with ice-cold extraction buffer containing 25 mM Tris-HCl (pH 7.5), 75 mM NaCl, 5% (v/v) glycerol, 0.05% (v/v) Nonidet P-40, 0.5 mM EDTA, 0.5 mM EGTA, 2 mM DTT, 2% (w/v) insoluble PVP (Sigma P-6755), supplemented with 1X Protease and Phosphatase inhibitor cocktail (Thermo-Fisher; cat# PIA32959). Total protein content was quantified by Bradford assay (Thermo-Fisher, cat# 23236).

For immunoblot analysis, 50 µg of total protein was separated on a 12.5% (w/v) SDS-PAGE gel and electroblotted onto polyvinylidene fluoride (PVDF) membrane. After 1 hr of blocking at 22 °C with TBST buffer (1X Tris-buffered saline [pH7.6], 0.1% Tween-20, 10% non-fat dry milk, and 0.2% BSA), the membrane was incubated overnight at 4 °C with polyclonal rabbit phospho-eIF2α antibody (Cell Signaling, cat # 9712S) diluted to 1:2000 in 1X TBST with 0.5% BSA. Following washing with 1X TBST, 10 min each for five repeats, the membrane was incubated with horseradish peroxidase conjugated anti-rabbit IgG (Vector labs, cat# PI-1000) for 1 hr at room temperature. After washing with 1X TBST, 10 min each for six repeats, horseradish peroxidase was detected using chemiluminescence (WesternBright Quantum, Advansta) as per manufacturer’s protocol. For immunoblot with polyclonal rabbit eIF2α antibody (a gift from Dr. Karen Browning, University of Texas, Austin), 5 µg of total protein was resolved by SDS-PAGE and electroblotted onto a polyvinylidene difluoride (PVDF) membrane. Blocking and incubation with antibodies was performed as described in (Dennis et al., 2009) followed by chemiluminescent detection (Lokdarshi et al., 2016).

### Polysome profiling and protein fractionation

Two-week-old seedlings were flash frozen in liquid N_2_ and stored at -80 °C prior to polysome profiling. Whole seedlings were ground in liquid N_2_ and 0.5g of tissue powder was resuspended in 1 mL of polysome extraction buffer (200 mM Tris-HCl pH 8.4, 50 mM KCl, 25 mM MgCl_2_, 1% deoxycholic acid, 2% polyoxyethylene 10 tridecyl ether, 50 µg/mL cycloheximide and 40U/mL RNase inhibitor (Promega Cat# N2115) and centrifuged at 13,000 x *g* for 5 min at 4 °C. One mL of the supernatant was layered onto a 10 mL 15-50% linear gradient prepared using a Hoefer gradient maker and centrifuged at 35,000 rpm (Beckmann SW 41 Ti) for 3.5 hr at 4 °C. Absorbance at 254 nm was recorded using an ISCO UA 5 absorbance/fluorescence monitor and individual data points were extracted using the DATA acquisition software (DATAQ instruments). Polysome-to-monosome (P/M) ratios were calculated as described (Enganti et al., 2018).

### Reactive oxygen species localization and microscopic techniques

For subcellular detection of ROS in Arabidopsis leaves, stress exposed 2-week-old seedlings were submerged in 15µM H_2_DCFDA (ThermoFisher, cat# D339) for 10-12 min in the dark. ROS detection was performed on a Leica SP8 laser scanning confocal microscope using the HeNe laser in the Advanced Microscopy and Imaging Facility at the University of Tennessee, Knoxville. The excitation filter was set to 488nm and the emission filter was to 500-550 nm for H_2_DCFDA (emission maximum of 517-527nm) and 660-690 nm for chlorophyll autofluorescence. Post processing of confocal z-stack images was performed using ImageJ (ver. 1.4; http://rsb.info.nih.gov/ij/index.html).

### Hydrogen peroxide quantification

H_2_O_2_ measurements were performed using the Amplex Red kit (Thermo-Fisher, cat# A22188). Briefly, 30 mg of 2-week-old seedling leaves were flash frozen in liquid N_2_ and ground with a plastic pestle to a homogeneous powder. Pulverized tissue was resuspended in 100 µl of sterile 1X phosphate buffered saline (PBS) and centrifuged at 17000 x g 4 °C for 2 minutes and the supernatant was used for H_2_O_2_ measurements as per manufacturer’s protocol. Relative fluorescence was measured on a POLARstar OPTIMA plate reader (BMG LABTECH) with an excitation filter at 535 nm and emission filter at 600 nm.

### Photosynthetic efficiency measurement

The maximum quantum yield of photosystem II [Qymax= F_v_ /F_m_] was measured on a FluorCam 800MF (Photon Systems Instruments) as per manufacturer’s instructions and modifications from (Murchie and Lawson, 2013). Briefly, 16 hr light/8 hr dark grown 3-day-old wild-type Landsberg and *gcn2-1* seedlings were dark adapted for 2 min (F_0_) prior to applying a saturating pulse of 1800 µEin m^-2^s^-1^ for 0.8 sec (F_m_). Variable fluorescence (F_v_) was calculated as the difference between F_o_ and F_m_ to get the maximum quantum yield [F_v_/F_m_].

### Transcriptome analysis

For microarray analysis, seedlings were grown as described (Missra and von Arnim, 2014; Missra et al., 2015). Eleven-day-old WT or *gcn2-1* seedlings were treated around ZT 2 by dipping in a 0.5 µM chlorosulfuron solution in water or in a mock treatment for 2 min. Treated seedling shoots were harvested 2 hours after treatment by freezing in liquid nitrogen and scraping off the aerial tissue with a metal spatula. Total RNA extraction and purification as well as Affymetrix GeneChip *Arabidopsis* ATH1 Genome Array hybridization and scanning were performed as described (Missra and von Arnim, 2014). Each treatment/genotype combination was performed in triplicate resulting in 4 treatment/genotype combinations and a total of 12 microarrays.

The microarrays were analyzed in R (ver. 3.5.0) (R Core Team, 2018) and normalized using the gcrma package (ver. 2.52.0) (Wu et al., 2018) with default parameters. The mas5calls function was used to create present, marginal or absent calls from the normalized data. Probes were filtered by mas5 calls, only keeping probes which expressed only present values in all three replicates of at least one condition. Probes for chloroplastic, mitochondrial, and control genes as well as probes with 0 variance across all 12 microarrays were also removed. Differentially expressed genes (DEGs) were identified by using Limma (version 3.36.2) (Ritchie et al., 2015; Phipson et al., 2016). DEGs were identified using a factorial design with terms included to account for a replicate batch effect in the data. DEG results were corrected for multiple comparisons using FDR. A one-way ANOVA using limma’s topTableF ftest was used to find genes that had equal means between conditions for future filtering, the p-values were corrected by the Benjamini-Hochberg method. This helped to define a set of non-differentially expressed genes.

Following differential expression analysis, the intersection of genes found to be significant in the one-way ANOVA and limma’s factorial design were then divided according to expression patterns. The first group contained genes with a fold change of less or equal to 1.3 or did not have an FDR < 0.05 and were considered not differentially expressed. A second group contained genes with a fold change greater than 1.3 and FDR < 0.05. This second group was further subdivided into 7 gene sets where differential gene expression occurred: 1) only in the WT; 2) only in the *gcn2* mutant; 3-5) upregulated or downregulated in both genotypes with subgroups 3) WT = *gcn2* (i.e. |WT-*gcn2*| < 1.3 fold change), 4) *gcn2* > WT and 5) WT > *gcn2*; 6) opposite differential expression in WT and *gcn2* (*e.g.* up in WT and down in *gcn2* or *vice versa*); and 7) differentially expressed in between WT and *gcn2* controls, but not in the herbicide-treated conditions. These seven subgroups were reduced to five by pooling closely related groups and subdivided again into sets containing only upregulated and downregulated genes (total of 11 groups: 1 no differential expression, 5 upregulated, 5 downregulated).

The replicate heat map was created with the hclust package in R using Pearson’s R-square as distances and the complete algorithm. In the heat map, the expression level (X_ijk_) of gene i, in replicate j under condition k is displayed as a Z-score with Z_ijk_ = (X_ijk_-AVERAGE_i_)/STDEV_i_. Gene ontology was performed on the 15 gene sets using the topGO package (version 2.32.0) (Alexa and Rahnenfuhrer, 2016) and revision 46221 (http://viewvc.geneontology.org/viewvc/GO-SVN/trunk/gene-associations/gene_association.tair.gz?revision=46221) of the TAIR gene association files from the gene ontology consortium. Only genes measured as expressed (15,934 genes) were used as the gene universe, topGO was run with node size 1, and FDR p-value adjustment using a custom script and the classic Fisher, parent-child and weight01 algorithms. Packages were obtained either from CRAN or from Bioconductor (version 3.7) (Huber et al., 2015).

### Translatome analysis

After sucrose gradient fractionation of cell extracts from seedling shoots, we generated pools of non-polysomal (NP), small polysomal (SP) and large polysomal (LP) RNAs. The RNAs were processed for ATH1 Affymetrix RNA hybridization to measure the abundance of each mRNA in each fraction. Hybridization signals were extracted and normalized using the standard gcRMA algorithm. Subsequently, a translation state (TL) was calculated for each mRNA, essentially as described in Missra et al. (2015) as follows; TL = (0*NP + 2*SP + 7*LP) / (NP + SP + LP). From the wild type dataset, one of the three replicates was eliminated because of excessive variance compared with the other two replicates; we assert that this variance is due to a technical flaw in the experiment. We initially did not adjust the data to account for the global loss of polysome loading in herbicide treated wild type. In a subsequent, alternate analysis, a scaling factor was applied to the array data from herbicide-treated wild type in order to mimic the global repression of translation that was evident after RNA isolation. Differentially translated mRNAs were identified by Limma with FDR correction (p < 0.05). These were clustered by hierarchical clustering with Pearson correlation, the preferred cluster number was settled upon by silhouette score analysis, and TL values were displayed as a Z-score (the fold difference from the average divided by the standard deviation across each of four treatments). Clusters were analyzed for functional enrichment using topGO. Pairwise comparisons were analyzed using the pre-ranked function in the gene set enrichment analysis suite (Mootha et al. 2003, Subramanian et al. 2005)

### Accession numbers

Arabidopsis AGI locus identifiers are At3G59410 (*GCN2*), and At5g05470/At2g40290 (*eIF2*α). Microarray data are at NCBI GEO accession GSE52117.

## Supplemental Data Files

**Supplemental Table S1. Translatome data gene ontology.**

**Supplemental Figure 1. Heat stress suppresses eIF2*α* phosphorylation.**

**Supplemental Figure 2. eIF2*α* phosphorylation under high light stress.**

**Supplemental Figure 3. Dark acclimation suppresses eIF2*α* phosphorylation.**

**Supplemental Figure 4. Ascorbate delays GCN2 activation in the light.**

**Supplementary Figure 5. Characterization of the *gcn2-2* and *gcn2-3* T-DNA insertion alleles of *GCN2*.**

**Supplemental Figure 6. The *gcn2-2* allele in Columbia ecotype shows increased sensitivity towards continuous light and high light treatments.**

**Supplemental Figure 7. The *gcn2* mutant alleles in the Columbia ecotype shows elevated sensitivity towards continuous or high light.**

**Supplemental Figure 8. GCN2 has minimal effects on photosystem II quantum yield (Fv/Fm).**

**Supplemental Figure 9. Dark to light shift modulates hydrogen peroxide levels.**

**Supplemental Figure 10. Hydrogen peroxide treatment represses translation similarly in *gcn2-1* mutant and wild-type.**

**Supplemental Figure 11. Norflurazon induces GCN2 activity in a manner dependent on photosynthesis.**

**Supplemental Figure 12. Activation of GCN2 by methyl viologen requires light.**

**Supplemental Figure 13. Effect of methyl viologen on photosynthetic efficiency of wild-type and *gcn2-1*.**

**Supplemental Figure 14. Treatments intended to induce ROS in mitochondria and ER cause accumulation of H**_**2**_**O**_**2**_.

**Supplemental Figure 15. *gcn2* mutant and wild-type show similar ribosome RNA profile upon dark adaptation and after return to light.**

**Supplemental Figure 16. GCN2 imparts bacterial resistance by chitin-inducible priming.**

**Supplemental Data Set 1. Translation state data. Supplemental Data Set 2. Transcriptome data.**

## ACKNOWLEDGMENTS

This work was supported by grants from the National Science Foundation (IOS-1456988 and NSF-MCB 1546402) to AGvA. AGvA acknowledges generous support through the Donald L Akers Jr Faculty Enrichment Fellowship. Funding in the laboratory of BD was supported by the National Science Foundation (IOS-1557437) and the National Institutes of General Medical Sciences (1R01GM125743). We thank Karen Browning for the gift of antibody against eIF2α. We also thank Dr. Chanhong Kim for sharing *flu1-1* seeds and Dr. Veronica G. Maurino for the glycolate oxidase (GO) mutants.

## AUTHOR CONTRIBUTIONS

Designed the research: AL, JG, SKC, RAUC, MS, BD, AGvA; performed research: AL, JG, SKC, RAUC, PM, ML, MS; analyzed data: AL, JG, SKC, RAUC, MS, BD, AGvA; wrote the paper: AL, BD, AGvA.

## REFERENCES

Alboresi, A., Dall’Osto, L., Aprile, A., Carillo, P., Roncaglia, E., Cattivelli, L., and Bassi, R. (2011). Reactive oxygen species and transcript analysis upon excess light treatment in wild-type Arabidopsis thaliana vs a photosensitive mutant lacking zeaxanthin and lutein. BMC Plant Biology 11, 62.

Alexa, A., and Rahnenfuhrer, J. (2016). topGO: Enrichment Analysis for Gene Ontology October 30, 2018. http://www.mpi-sb.mpg.de/alexa

Alonso, J.M., Stepanova, A.N., Leisse, T.J., Kim, C.J., Chen, H., Shinn, P., Stevenson, D.K., Zimmerman, J., Barajas, P., Cheuk, R., Gadrinab, C., Heller, C., Jeske, A., Koesema, E., Meyers, C.C., Parker, H., Prednis, L., Ansari, Y., Choy, N., Deen, H., Geralt, M., Hazari, N., Hom, E., Karnes, M., Mulholland, C., Ndubaku, R., Schmidt, I., Guzman, P., Aguilar-Henonin, L., Schmid, M., Weigel, D., Carter, D.E., Marchand, T., Risseeuw, E., Brogden, D., Zeko, A., Crosby, W.L., Berry, C.C., and Ecker, J.R. (2003). Genome-wide insertional mutagenesis of Arabidopsis thaliana. Science 301, 653–657.

Anda, S., Zach, R., and Grallert, B. (2017). Activation of Gcn2 in response to different stresses. PLoS One 12, e0182143.

Akbudak, N., Tezcan, H., Akbudak, B., and Seniz, V. (2006). The effect of harpin protein on plant growth parameters, leaf chlorophyll, leaf colour and percentage rotten fruit of pepper plants inoculated with Botrytis cinerea. Scientia Horticulturae 109, 107–112.

Asada, K. (1999). The water-water cycle in chloroplasts: Scavenging of active oxygens and dissipation of excess photons. Annu Rev Plant Physiol Plant Mol Biol 50, 601–639.

Asada, K. (2006). Production and scavenging of reactive oxygen species in chloroplasts and their functions. Plant Physiology 141, 391–396.

Baxter, A., Mittler, R., and Suzuki, N. (2014). ROS as key players in plant stress signalling. J Exp Bot 65, 1229–1240.

Berlanga, J.J., Ventoso, I., Harding, H.P., Deng, J., Ron, D., Sonenberg, N., Carrasco, L., and de Haro, C. (2006). Antiviral effect of the mammalian translation initiation factor 2*α* kinase GCN2 against RNA viruses. Embo J 25, 1730–1740.

Bode, R., Ivanov, A.G., and Huner, N.P. (2016). Global transcriptome analyses provide evidence that chloroplast redox state contributes to intracellular as well as long-distance signalling in response to stress and acclimation in Arabidopsis. Photosynthesis Research 128, 287–312.

Camejo, D., Guzman-Cedeno, A., and Moreno, A. (2016). Reactive oxygen species, essential molecules, during plant-pathogen interactions. Plant Physiol Biochem 103, 10–23.

Cao, S.S., and Kaufman, R.J. (2014). Endoplasmic reticulum stress and oxidative stress in cell fate decision and human disease. Antioxid Redox Sign 21, 396–413.

Chakrabarti, S., Liehl, P., Buchon, N., and Lemaitre, B. (2012). Infection-induced host translational blockage inhibits immune responses and epithelial renewal in the Drosophila gut. Cell Host Microbe 12, 60–70.

Choudhury, F.K., Rivero, R.M., Blumwald, E., and Mittler, R. (2017). Reactive oxygen species, abiotic stress and stress combination. Plant J 90, 856–867.

Crisp, P.A., Ganguly, D.R., Smith, A.B., Murray, K.D., Estavillo, G.M., Searle, I., Ford, E., Bogdanovic, O., Lister, R., Borevitz, J.O., Eichten, S.R., and Pogson, B.J. (2017). Rapid Recovery Gene Downregulation during Excess-Light Stress and Recovery in Arabidopsis. Plant Cell 29, 1836–1863.

Cui, F., Brosche, M., Shapiguzov, A., He, X.-Q., Vainonen, J., Leppälä, J., Trotta, A., Kangasjärvi, S., Salojärvi, J., Kangasjärvi, J., and Overmyer, K. (2018). Methyl viologen can affect mitochondrial function in Arabidopsis. bioRxiv Oct 7, 2018. doi: https://doi.org/10.1101/436543

Czarnocka, W., and Karpinski, S. (2018). Friend or foe? Reactive oxygen species production, scavenging and signaling in plant response to environmental stresses. Free Radic Biol Med 122, 4–20.

Das, K., and Roychoudhury, A. (2014). Reactive oxygen species (ROS) and response of antioxidants as ROS-scavengers during environmental stress in plants. Frontiers in Environmental Science 2, 53.

Del Rio, L.A., and Lopez-Huertas, E. (2016). ROS Generation in Peroxisomes and its Role in Cell Signaling. Plant Cell Physiol 57, 1364–1376.

Demidchik, V. (2015). Mechanisms of oxidative stress in plants: From classical chemistry to cell biology. Environ Exp Bot 109, 212–228.

Dennis, M.D., Person, M.D., and Browning, K.S. (2009). Phosphorylation of plant translation initiation factors by CK2 enhances the in vitro interaction of multifactor complex components. J Biol Chem 284, 20615–20628.

Dever, T.E., Feng, L., Wek, R.C., Cigan, A.M., Donahue, T.F., and Hinnebusch, A.G. (1992). Phosphorylation of initiation factor 2*α* by protein kinase GCN2 mediates gene-specific translational control of GCN4 in yeast. Cell 68, 585–596.

Dietz, K.J. (2015). Efficient high light acclimation involves rapid processes at multiple mechanistic levels. J Exp Bot 66, 2401–2414.

Dietz, K.J., Turkan, I., and Krieger-Liszkay, A. (2016). Redox- and Reactive Oxygen Species-Dependent Signaling into and out of the Photosynthesizing Chloroplast. Plant Physiol 171, 1541–1550.

Dong, J.S., Qiu, H.F., Garcia-Barrio, M., Anderson, J., and Hinnebusch, A.G. (2000). Uncharged tRNA activates GCN2 by displacing the protein kinase moiety from a bipartite tRNA-Binding domain. Mol Cell 6, 269–279.

Dong, Y., Silbermann, M., Speiser, A., Forieri, I., Linster, E., Poschet, G., Allboje Samami, A., Wanatabe, M., Sticht, C., Teleman, A.A., Deragon, J.M., Saito, K., Hell, R., and Wirtz, M. (2017). Sulfur availability regulates plant growth via glucose-TOR signaling. Nat Commun 8, 1174.

Duncan, C.D.S., Rodriguez-Lopez, M., Ruis, P., Bahler, J., and Mata, J. (2018). General amino acid control in fission yeast is regulated by a nonconserved transcription factor, with functions analogous to Gcn4/Atf4. P Natl Acad Sci USA 115, E1829–E1838.

Ehonen, S., Yarmolinsky, D., Kollist, H., and Kangasjarvi, J. (2018). Reactive Oxygen Species, Photosynthesis, and Environment in the Regulation of Stomata. Antioxid Redox Sign doi: 10.1089/ars.2017.7455.

Enganti, R., Cho, S.K., Toperzer, J.D., Urquidi-Camacho, R.A., Cakir, O.S., Ray, A.P., Abraham, P.E., Hettich, R.L., and von Arnim, A.G. (2018). Phosphorylation of Ribosomal Protein RPS6 Integrates Light Signals and Circadian Clock Signals. Front Plant Sci 8.

Fahnenstich, H., Scarpeci, T.E., Valle, E.M., Flügge, U.I., and Maurino, V.G. (2008). Generation of hydrogen peroxide in chloroplasts of Arabidopsis overexpressing glycolate oxidase as an inducible system to study oxidative stress. Plant Physiol 148, 719–729.

Faus, I., Zabalza, A., Santiago, J., Nebauer, S.G., Royuela, M., Serrano, R., and Gadea, J. (2015). Protein kinase GCN2 mediates responses to glyphosate in Arabidopsis. BMC Plant Biol 15, 14.

Foyer, C.H., and Noctor, G. (2016). Stress-triggered redox signalling: what’s in pROSpect? Plant Cell Environ 39, 951–964.

Fuerst, E.P., and Norman, M.A. (1991). Interactions of Herbicides with Photosynthetic Electron-Transport. Weed Sci 39, 458–464.

Fujii, T., Yokoyama, E., Inoue, K., and Sakurai, H. (1990). The Sites of Electron Donation of Photosystem-I to Methyl Viologen. Biochimica et Biophysica Acta 1015, 41–48.

Galvez-Valdivieso, G., Fryer, M.J., Lawson, T., Slattery, K., Truman, W., Smirnoff, N., Asami, T., Davies, W.J., Jones, A.M., Baker, N.R., and Mullineaux, P.M. (2009). The high light response in Arabidopsis involves ABA signaling between vascular and bundle sheath cells. Plant Cell 21, 2143–2162.

Huber, W., Carey, V.J., Gentleman, R., Anders, S., Carlson, M., Carvalho, B.S., Bravo, H.C., Davis, S., Gatto, L., Girke, T., Gottardo, R., Hahne, F., Hansen, K.D., Irizarry, R.A., Lawrence, M., Love, M.I., MacDonald, J., Obenchain, V., Oles, A.K., Pages, H., Reyes, A., Shannon, P., Smyth, G.K., Tenenbaum, D., Waldron, L., and Morgan, M. (2015). Orchestrating high-throughput genomic analysis with Bioconductor. Nat Methods 12, 115–121.

Izquierdo, Y., Kulasekaran, S., Benito, P., Lopez, B., Marcos, R., Cascon, T., Hamberg, M., and Castresana, C. (2018). Arabidopsis nonresponding to oxylipins locus NOXY7 encodes a yeast GCN1 homolog that mediates noncanonical translation regulation and stress adaptation. Plant Cell and Environment 41, 1438–1452.

Jacques, S., Ghesquiere, B., Van Breusegem, F., and Gevaert, K. (2013). Plant proteins under oxidative attack. Proteomics 13, 932–940.

Jaipargas, E.A., Mathur, N., Bou Daher, F., Wasteneys, G.O., and Mathur, J. (2016). High Light Intensity Leads to Increased Peroxule-Mitochondria Interactions in Plants. Front Cell Dev Biol 4, 6.

Juntawong, P., and Bailey-Serres, J. (2012). Dynamic light regulation of translation status in Arabidopsis thaliana. Front Plant Sci 3, 66.

Khandal, D., Samol, I., Buhr, F., Pollmann, S., Schmidt, H., Clemens, S., Reinbothe, S., and Reinbothe, C. (2009). Singlet oxygen-dependent translational control in the tigrina-d.12 mutant of barley. Proc Natl Acad Sci USA 106, 13112–13117.

Khorobrykh, S.A., Karonen, M., and Tyystjärvi, E. (2015). Experimental evidence suggesting that H2O2 is produced within the thylakoid membrane in a reaction between plastoquinol and singlet oxygen. FEBS Letters 589, 779–786.

Knutsen, J.H., Rodland, G.E., Boe, C.A., Haland, T.W., Sunnerhagen, P., Grallert, B., and Boye, E. (2015). Stress-induced inhibition of translation independently of eIF2alpha phosphorylation. J Cell Sci 128, 4420–4427.

Kruk, J., and Karpinski, S. (2006). An HPLC-based method of estimation of the total redox state of plastoquinone in chloroplasts, the size of the photochemically active plastoquinone-pool and its redox state in thylakoids of Arabidopsis. BBA-Bioenergetics 1757, 1669–1675.

Lageix, S., Lanet, E., Pouch-Pelissier, M.N., Espagnol, M.C., Robaglia, C., Deragon, J.M., and Pelissier, T. (2008). Arabidopsis eIF2*α* kinase GCN2 is essential for growth in stress conditions and is activated by wounding. BMC Plant Biol 8, 134.

Lai, A.G., Doherty, C.J., Mueller-Roeber, B., Kay, S.A., Schippers, J.H., and Dijkwel, P.P. (2012). CIRCADIAN CLOCK-ASSOCIATED 1 regulates ROS homeostasis and oxidative stress responses. Proc Natl Acad Sci U S A 109, 17129–17134.

Leitman, J., Barak, B., Benyair, R., Shenkman, M., Ashery, U., Hartl, F.U., and Lederkremer, G.Z. (2014). ER Stress-Induced eIF2-*α* Phosphorylation Underlies Sensitivity of Striatal Neurons to Pathogenic Huntingtin. Plos One 9, e90803.

Li, M.W., AuYeung, W.K., and Lam, H.M. (2013). The GCN2 homologue in Arabidopsis thaliana interacts with uncharged tRNA and uses Arabidopsis eIF2*α* molecules as direct substrates. Plant Biol (Stuttg) 15, 13–18.

Li, Z., Wakao, S., Fischer, B.B., and Niyogi, K.K. (2009). Sensing and responding to excess light. Annu Rev Plant Biol 60, 239–260.

Liu, M.J., Wu, S.H., Chen, H.M., and Wu, S.H. (2012). Widespread translational control contributes to the regulation of Arabidopsis photomorphogenesis. Mol Syst Biol 8, 566.

Liu, X., Afrin, T., and Pajerowska-Mukhtar, K.M. (2019). Arabidopsis GCN2 kinase contributes to ABA homeostasis and stomatal immunity. Commun Biol 2, 302.

Liu, X., Merchant, A., Rockett, K.S., McCormack, M., and Pajerowska-Mukhtar, K.M. (2015). Characterization of Arabidopsis thaliana GCN2 kinase roles in seed germination and plant development. Plant Signal Behav 10, e992264.

Lokdarshi, A., Conner, W.C., McClintock, C., Li, T., and Roberts, D.M. (2016). Arabidopsis CML38, a Calcium Sensor That Localizes to Ribonucleoprotein Complexes under Hypoxia Stress. Plant Physiol 170, 1046–1059.

Luna, E., van Hulten, M., Zhang, Y., Berkowitz, O., Lopez, A., Petriacq, P., Sellwood, M.A., Chen, B., Burrell, M., van de Meene, A., Pieterse, C.M., Flors, V., and Ton, J. (2014). Plant perception of beta-aminobutyric acid is mediated by an aspartyl-tRNA synthetase. Nat Chem Biol 10, 450–456.

Mateo, A., Muhlenbock, P., Rusterucci, C., Chang, C.C., Miszalski, Z., Karpinska, B., Parker, J.E., Mullineaux, P.M., and Karpinski, S. (2004). Lesion simulating disease 1 - Is required for acclimation to conditions that promote excess excitation energy. Plant Physiology 136, 2818–2830.

Maxwell, D.P., Wang, Y., and McIntosh, L. (1999). The alternative oxidase lowers mitochondrial reactive oxygen production in plant cells. Proc Natl Acad Sci U S A 96, 8271–8276.

Merchante, C., Stepanova, A.N., and Alonso, J.M. (2017). Translation regulation in plants: an interesting past, an exciting present and a promising future. Plant J 90, 628–653.

Meskauskiene, R., Nater, M., Goslings, D., Kessler, F., op den Camp, R., and Apel, K. (2001). FLU: a negative regulator of chlorophyll biosynthesis in Arabidopsis thaliana. Proc Natl Acad Sci U S A 98, 12826–12831.

Mignolet-Spruyt, L., Xu, E., Idanheimo, N., Hoeberichts, F.A., Muhlenbock, P., Brosche, M., Van Breusegem, F., and Kangasjarvi, J. (2016). Spreading the news: subcellular and organellar reactive oxygen species production and signalling. J Exp Bot 67, 3831–3844.

Missra, A., and von Arnim, A.G. (2014). Analysis of mRNA translation states in Arabidopsis over the diurnal cycle by polysome microarray. Methods Mol Biol 1158, 157–174.

Missra, A., Ernest, B., Lohoff, T., Jia, Q., Satterlee, J., Ke, K., and von Arnim, A.G. (2015). The Circadian Clock Modulates Global Daily Cycles of mRNA Ribosome Loading. Plant Cell 27, 2582–2599.

Mittler, R. (2017). ROS Are Good. Trends Plant Sci 22, 11–19.

Moore, M., Gossmann, N., and Dietz, K.J. (2016). Redox Regulation of Cytosolic Translation in Plants. Trends Plant Sci 21, 388–397.

Mootha, V.K., Lindgren, C.M., Eriksson, K.F., Subramanian, A., Sihag, S., Lehar, J., Puigserver, P., Carlsson, E., Ridderstråle, M., Laurila, E., Houstis, N., Daly, MJ., Patterson, N., Mesirov, JP., Golub, T.R., Tamayo, P., Spiegelman, B., Lander, E.S., Hirschhorn, J.N., Altshuler, D., Groop, L.C. (2003) PGC-1alpha-responsive genes involved in oxidative phosphorylation are coordinately downregulated in human diabetes. Nat Genet 34, 267–73.

Mubarakshina, M.M., and Ivanov, B.N. (2010). The production and scavenging of reactive oxygen species in the plastoquinone pool of chloroplast thylakoid membranes. Physiol Plantarum 140, 103–110.

Mullineaux, P.M., Exposito-Rodriguez, M., Laissue, P.P., and Smirnoff, N. (2018). ROS-dependent signalling pathways in plants and algae exposed to high light: Comparisons with other eukaryotes. Free Radic Biol Med 122, 52–64.

Munoz, P., and Munne-Bosch, S. (2018). Photo-Oxidative Stress during Leaf, Flower and Fruit Development. Plant Physiology 176, 1004–1014.

Murchie, E.H., and Lawson, T. (2013). Chlorophyll fluorescence analysis: a guide to good practice and understanding some new applications. J Exp Bot 64, 3983–3998.

op den Camp, R.G., Przybyla, D., Ochsenbein, C., Laloi, C., Kim, C., Danon, A., Wagner, D., Hideg, E., Gobel, C., Feussner, I., Nater, M., and Apel, K. (2003). Rapid induction of distinct stress responses after the release of singlet oxygen in Arabidopsis. Plant Cell 15, 2320–2332.

Ozgur, R., Turkan, I., Uzilday, B., and Sekmen, A.H. (2014). Endoplasmic reticulum stress triggers ROS signalling, changes the redox state, and regulates the antioxidant defence of Arabidopsis thaliana. J Exp Bot 65, 1377–1390.

Paradiso, A., Caretto, S., Leone, A., Bove, A., Nisi, R., and De Gara, L. (2016). ROS Production and Scavenging under Anoxia and Re-Oxygenation in Arabidopsis Cells: A Balance between Redox Signaling and Impairment. Front Plant Sci 7, 1803.

Pei, Z.M., Murata, Y., Benning, G., Thomine, S., Klusener, B., Allen, G.J., Grill, E., and Schroeder, J.I. (2000). Calcium channels activated by hydrogen peroxide mediate abscisic acid signalling in guard cells. Nature 406, 731–734.

Phipson, B., Lee, S., Majewski, I.J., Alexander, W.S., and Smyth, G.K. (2016). Robust Hyperparameter Estimation Protects against Hypervariable Genes and Improves Power to Detect Differential Expression. Ann Appl Stat 10, 946–963.

R Core Team. (2018). R: A Language and Environment for Statistical Computing (Vienna, Austria: R Foundation for Statistical Computing).

Reinbothe, S., Reinbothe, C., and Parthier, B. (1993). Methyl jasmonate-regulated translation of nuclear-encoded chloroplast proteins in barley (Hordeum vulgare L. cv. salome). J Biol Chem 268, 10606–10611.

Ritchie, M.E., Phipson, B., Wu, D., Hu, Y., Law, C.W., Shi, W., and Smyth, G.K. (2015). limma powers differential expression analyses for RNA-sequencing and microarray studies. Nucleic Acids Res 43, e47.

Sandmann, G. (1994). Carotenoid biosynthesis in microorganisms and plants. Eur J Biochem 223, 7–24.

Schmitt, F.J., Renger, G., Friedrich, T., Kreslavski, V.D., Zharmukhamedov, S.K., Los, D.A., Kuznetsov, V.V., and Allakhverdiev, S.I. (2014). Reactive oxygen species: re-evaluation of generation, monitoring and role in stress-signaling in phototrophic organisms. Biochim Biophys Acta 1837, 835–848.

Sesma, A., Castresana, C., and Castellano, M.M. (2017). Regulation of Translation by TOR, eIF4E and eIF2*α* in Plants: Current Knowledge, Challenges and Future Perspectives. Front Plant Sci 8, 644.

Stonebloom, S., Brunkard, J.O., Cheung, A.C., Jiang, K., Feldman, L., and Zambryski, P. (2012). Redox states of plastids and mitochondria differentially regulate intercellular transport via plasmodesmata. Plant Physiol 158, 190–199.

Subramanian, A., Tamayo, P., Mootha, V.K., Mukherjee, S., Ebert, B.L., Gillette, M.A., Paulovich, A., Pomeroy, S.L., Golub, T.R., Lander, E.S., Mesirov J.P. (2005) Gene set enrichment analysis: a knowledge-based approach for interpreting genome-wide expression profiles. Proc Natl Acad Sci U S A 102, 15545–15550.

Takano, H.K., Beffa, R., Preston, C., Westra, P., and Dayan, F.E. (2019). Reactive oxygen species trigger the fast action of glufosinate. Planta 249, 1837–1849.

Tang, L., Bhat, S., and Petracek, M.E. (2003). Light control of nuclear gene mRNA abundance and translation in tobacco. Plant Physiol 133, 1979–1990.

Tripathy, B.C., and Oelmuller, R. (2012). Reactive oxygen species generation and signaling in plants. Plant Signal Behav 7, 1621–1633.

Tsukagoshi, H. (2016). Control of root growth and development by reactive oxygen species. Curr Opin Plant Biol 29, 57–63.

Vaahtera, L., Brosche, M., Wrzaczek, M., and Kangasjarvi, J. (2014). Specificity in ROS signaling and transcript signatures. Antioxid Redox Signal 21, 1422–1441.

Vanderauwera, S., Zimmermann, P., Rombauts, S., Vandenabeele, S., Langebartels, C., Gruissem, W., Inze, D., and Van Breusegem, F. (2005). Genome-wide analysis of hydrogen peroxide-regulated gene expression in Arabidopsis reveals a high light-induced transcriptional cluster involved in anthocyanin biosynthesis. Plant Physiol 139, 806–821.

Vetoshkina, D.V., Ivanov, B.N., Khorobrykh, S.A., Proskuryakov, I.I., and Borisova-Mubarakshina, M.M. (2017). Involvement of the chloroplast plastoquinone pool in the Mehler reaction. Physiol Plantarum 161, 45–55.

von Arnim, A.G., Jia, Q., and Vaughn, J.N. (2014). Regulation of plant translation by upstream open reading frames. Plant Sci 214, 1–12.

Waszczak, C., Carmody, M., and Kangasjärvi, J. (2018). Reactive Oxygen Species in Plant Signaling 69, 209–236.

Wek, R.C. (2018). Role of eIF2*α* Kinases in Translational Control and Adaptation to Cellular Stress. Cold Spring Harb Perspect Biol 10, pii: a032870.

Wek, R.C., Jackson, B.M., and Hinnebusch, A.G. (1989). Juxtaposition of domains homologous to protein kinases and histidyl-tRNA synthetases in GCN2 protein suggests a mechanism for coupling GCN4 expression to amino acid availability. Proc Natl Acad Sci U S A 86, 4579–4583.

Wek, S.A., Zhu, S., and Wek, R.C. (1995). The histidyl-tRNA synthetase-related sequence in the eIF-2*α* protein kinase GCN2 interacts with tRNA and is required for activation in response to starvation for different amino acids. Mol Cell Biol 15, 4497–4506.

Wu, J., Irizarry, R., MacDonald, J., and Gentry, J. (2018). gcrma: Background Adjustment Using Sequence Information. https://rdrr.io/bioc/gcrma/

Xu, J., Tran, T., Padilla Marcia, C.S., Braun, D.M., and Goggin, F.L. (2017). Superoxide-responsive gene expression in Arabidopsis thaliana and Zea mays. Plant Physiol Biochem 117, 51–60.

Zhan, K., Narasimhan, J., and Wek, R.C. (2004). Differential activation of eIF2 kinases in response to cellular stresses in Schizosaccharomyces pombe. Genetics 168, 1867–1875.

Zhang, Y., Dickinson, J.R., Paul, M.J., and Halford, N.G. (2003). Molecular cloning of an Arabidopsis homologue of GCN2, a protein kinase involved in co–ordinated response to amino acid starvation. Planta 217, 668–675.

Zhang, Y., Wang, Y., Kanyuka, K., Parry, M.A., Powers, S.J., and Halford, N.G. (2008). GCN2-dependent phosphorylation of eukaryotic translation initiation factor-2*α* in Arabidopsis. J Exp Bot 59, 3131–3141.

Zipfel, C., and Robatzek, S. (2010). Pathogen-associated molecular pattern-triggered immunity: veni, vidi…? Plant Physiol 154, 551–554.

