## Supplemental Figure 1 for "Light activates the translational regulatory GCN2 kinase via reactive oxygen species emanating from the chloroplast"

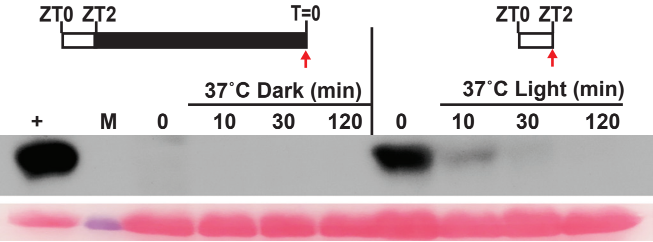


**Supplemental Figure 1. Heat stress suppresses eIF2$\alpha$ phosphorylation.** Supports Figure 1.
Time course of eIF2$\alpha$ phosphorylation in 14-days-old wild-type Landsberg seedlings in response to heat stress (37°C) after 24 hr dark acclimation (Dark) or after an additional 2 hr into the light period (Light).
The red arrow indicates the 0 minute time point. The lower panel shows the Rubisco large subunit (~55kDa) in the total protein sample after Ponceau S staining as a loading control. (+) arbitrary amount of total protein extract from glyphosate treated Wt seedlings indicating phosphorylated (eIF2$\alpha$-P) protein (~38kDa); (10, 30, 120) sampling time in minutes; (M) Molecular weight marker.
