## Supplemental Figure 2 for "Light activates the translational regulatory GCN2 kinase via reactive oxygen species emanating from the chloroplast"

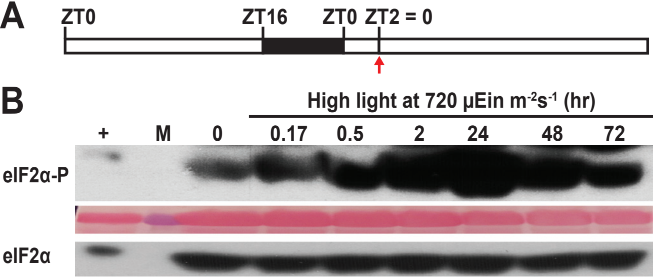


**Supplemental Figure 2. eIF2$\alpha$ phosphorylation under high light stress.** Supports Figure 1, 6.

**(A)** Wild-type Landsberg seedlings were grown for 3 days under a 16 hr light 8 hr dark period and shifted to high light at 780 µEin $m^{-2}s^{-1}$ starting at ZT2.

**(B)** Time course of eIF2$\alpha$ phosphorylation over 72 hr. Total eIF2$\alpha$ levels remained constant, as indicated. For details see legend to Supplemental Figure 1.
