## Supplemental Figure 3 for "Light activates the translational regulatory GCN2 kinase via reactive oxygen species emanating from the chloroplast"

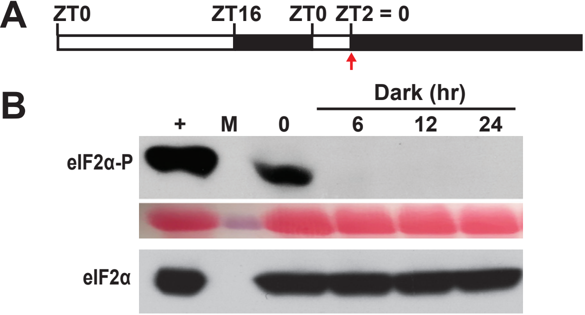


**Supplemental Figure 3. Dark acclimation suppresses eIF2$\alpha$ phosphorylation.** Supports Figure 1-4, 7-9.
**(A)** Schematic of the light regimen.
**(B)** Time course of eIF2$\alpha$ phosphorylation in 14-days-old wild-type Landsberg seedlings during dark acclimation (Dark). For details see legend to Supplemental Figure 1.
