## Supplemental Figure 4 for "Light activates the translational regulatory GCN2 kinase via reactive oxygen species emanating from the chloroplast"

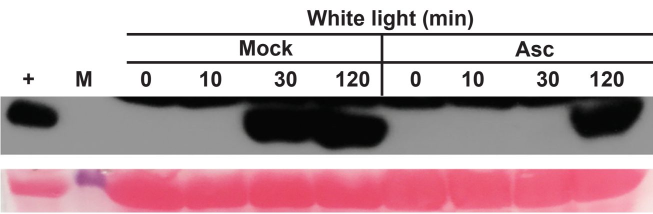


**Supplemental Figure 4. Ascorbate delays GCN2 activation in the light**. Supports Figure 4.
Time course of eIF2$\alpha$ phosphorylation in 14-days-old wild-type Landsberg seedlings grown on media supplemented with 0.5mM ascorbate (Asc). Seedlings were 24 hr dark acclimated and then exposed to light as described in Fig. 1A.
