## Supplemental Figure 5 for "Light activates the translational regulatory GCN2 kinase via reactive oxygen species emanating from the chloroplast"

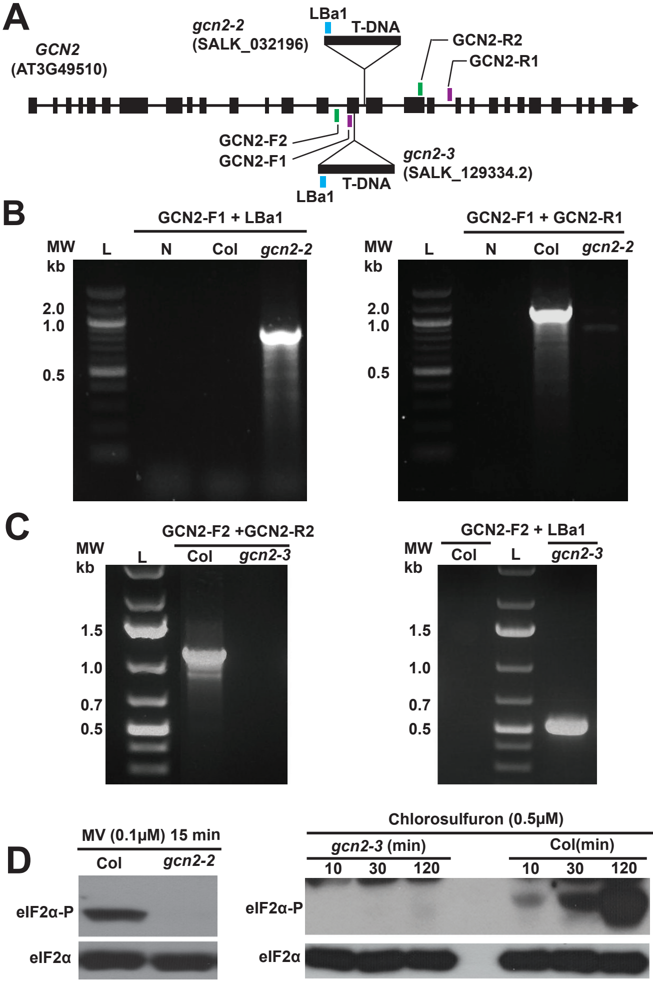


**Supplemental Figure 5. Characterization of the *gcn2-2* and *gcn2-3* T-DNA insertion alleles of *GCN2***. Supports Supplemental Figure 6,7.
**(A)** Schematic of the *GCN2* (AT3G49510) showing T-DNA insertion site in the 15th intron (4563bp downstream of the transcriptional start site) of *gcn2-2* and in the 15th exon of *gcn2-3* (4174bp downstream of the transcriptional start site). Colored bars indicate position of the primers used for PCR genotyping.
**(B,C)** Ethidium bromide stained 1% agarose (w/v) gel showing PCR-based genotyping of wild-type Columbia (Col), *gcn2-2* and *gcn2-3*. L, DNA molecular weight ladder (kb=kilo bases); N, no genomic DNA control. **(B)** Left panel - GCN2-F1+LBa1 (T-DNA left border primer), 892bp PCR product. Right panel - GCN2-F1+ GCN2-R1 (*GCN2* gene specific forward and reverse primer)**,** 1131bp PCR product. **(C)** Left panel - GCN2-F2+ GCN2-R2 (*GCN2* gene specific forward and reverse primer)**,** 1110bp PCR product. Right panel - GCN2-F2+LBa1, 520bp PCR product.
**(D)** Evidence that *gcn2-2* and *gcn2-3* behave as *gcn2* null allele. Seedlings were treated with indicated herbicides and probed for eIF2$\alpha$ phosphorylation. For details see legend to Fig 1.
