## Supplemental Figure 6 for "Light activates the translational regulatory GCN2 kinase via reactive oxygen species emanating from the chloroplast"

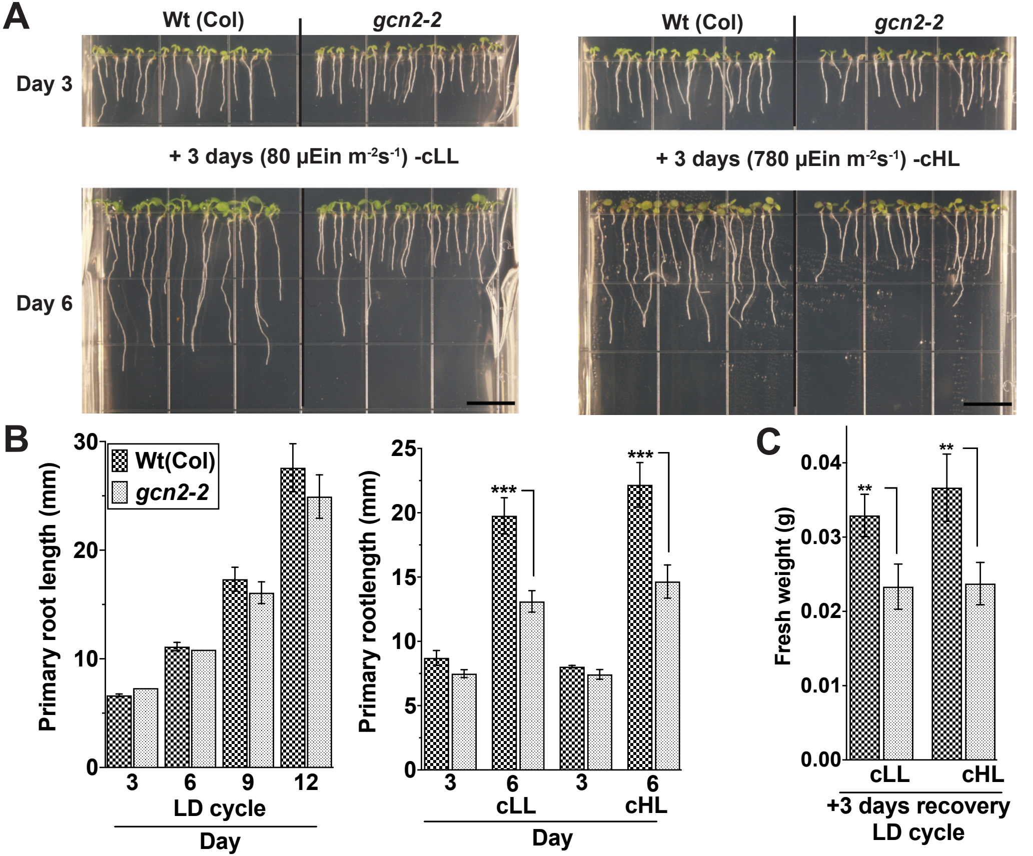


**Supplemental Figure 6. The *gcn2-2* allele in the Columbia ecotype shows increased sensitivity towards continuous light and high light treatments.** Supports Figure 6.

**(A)** Top panel, 3-days-old wild-type Columbia (Wt (Col)) and *gcn2-2* mutant *(gcn2-2)* seedlings grown under long day period (16 hr light, 8 hr dark). Bottom panel, Wt and *gcn2-2* mutants after 3 days of additional continuous light at 80 µEin $m^{-2}s^{-1}$ (cLL, left panel), or continuous high light at 780 µEin $m^{-2}s^{-1}$ (cHL, right panel). Scale bar, 10mm.

**(B)** Left - Primary root length of Wt and *gcn2-2* mutants grown under 16 hr light and 8 hr dark (LD) cycle, Right - Root length of Wt and *gcn2-2* as shown in panel (A).

**(C)** Seedling fresh weight of Wt and *gcn2-2* mutants after an additional 3 days of recovery after the cLL or cHL treatment. Error bars represent standard deviation of four biological replicates with n>12 (B-left) or n>80 (B-right, C) per experiment. (Welch's *t*-test ** P-value <0.005, *** P-value <0.0005).
