## Supplemental Figure 7 for "Light activates the translational regulatory GCN2 kinase via reactive oxygen species emanating from the chloroplast"

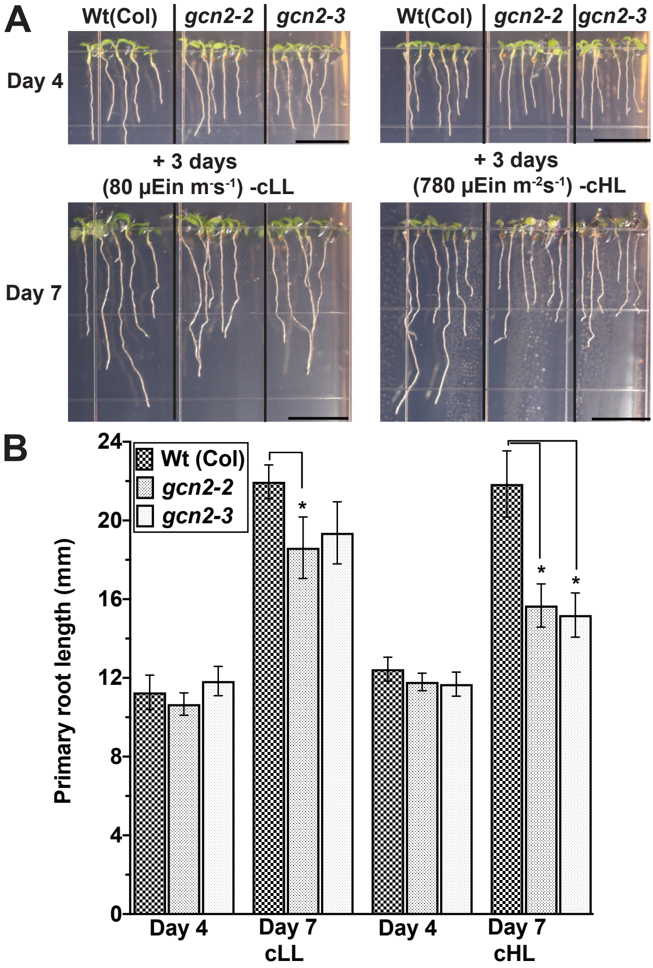


**Supplemental Figure 7. The *gcn2* mutant alleles in the Columbia ecotype show elevated sensitivity towards continuous or high light.**
Supports Figure 6, Supplemental Figure 6.
**(A)** Top panels - Wild-type Columbia (Wt (Col)) and *gcn2* T-DNA insertions *(gcn2-2, gcn2-3*) grown under long day period for 4-days. Bottom panels- Wt and *gcn2* mutants after 3 days of additional continuous light at 80 µEin $m^{-2}s^{-1}$ (cLL, left panel), or continuous high light at 780 µEin $m^{-2}s^{-1}$ (cHL, right panel). Scale bars are 10mm.
**(B)** Primary root length of Wt and *gcn2* mutants shown in panel (A). Error bars represent standard error of the mean from five technical replicates with n>25 (Welch's *t*-test *P-value <0.05).
