## Supplemental Figure 8 for "Light activates the translational regulatory GCN2 kinase via reactive oxygen species emanating from the chloroplast"

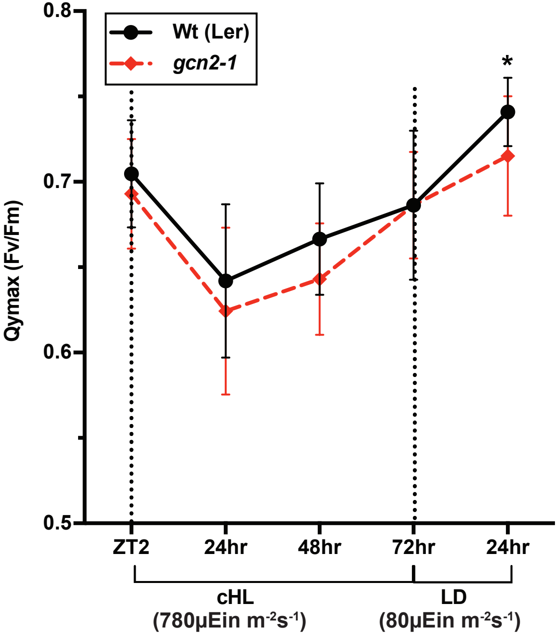


**Supplemental Figure 8. GCN2 has minimal effects on photosystem II quantum yield (Fv/Fm).** Supports Figure 6.
Twelve-days-old wild-type Landsberg (Wt (Ler)) and *gcn2-1* mutant *(gcn2-1)* seedlings were subjected to continuous high light stress (cHL (780μEin $m^{-2}s^{-1}$)) for 72 hr followed by 24 hr recovery at (LD (80μEin $m^{-2}s^{-1}$)) as indicated by the dashed lines. Error bars represent standard deviation of fifteen biological replicates. Welch's *t*-test, *P-value <0.05 .
