## Supplemental Figure 9 for "Light activates the translational regulatory GCN2 kinase via reactive oxygen species emanating from the chloroplast"

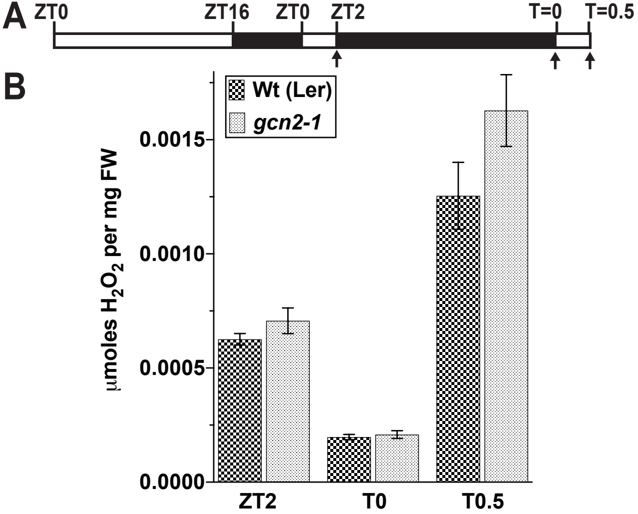


**Supplemental Figure 9. Dark to light shift modulates hydrogen peroxide levels.** Supports Figure 1, 6.

Relative $H_2O_2$ levels in 14-days-old wild-type Landsberg (Wt (Ler)) and *gcn2-1* mutant *(gcn2-1)* seedlings grown under a 16 hr light and 8 hr dark cycle. $H_2O_2$ was measured at 2 hr into the light (ZT2), after 24 hr dark acclimation (T0) and after another 30 minutes of 80 µEin $m^{-2}s^{-1}$ light exposure (T0.5). Error bars represent standard error of the mean of 8 biological replicates. Welch's *t*-test *P*-values were >0.5.
