## Supplemental Figure 10 for "Light activates the translational regulatory GCN2 kinase via reactive oxygen species emanating from the chloroplast"

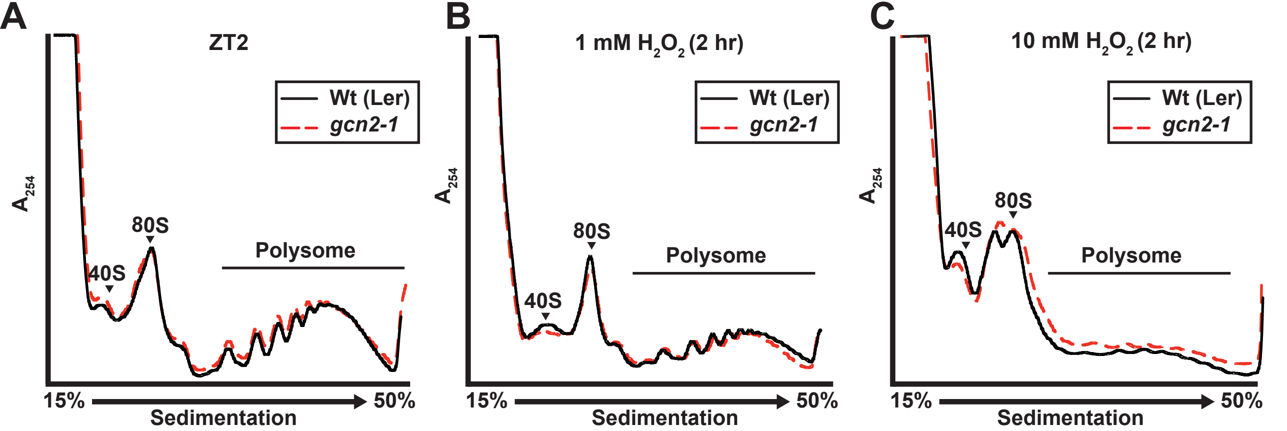


**Supplemental Figure 10. Hydrogen peroxide treatment represses translation similarly in *gcn2-1* mutant and wild-type.**
Supports Figure 6.
Seedlings grown in a long day period were sprayed with hydrogen peroxide ($H_2O_2$) at the indicated concentrations 2 hr after lights-on. Absorbance profiles (A254nm) at **(A)** ZT2, **(B)** after treatment with 1 mM $H_2O_2$ and, **(C)** 10 mM $H_2O_2$.
