## Supplemental Figure 11 for "Light activates the translational regulatory GCN2 kinase via reactive oxygen species emanating from the chloroplast"

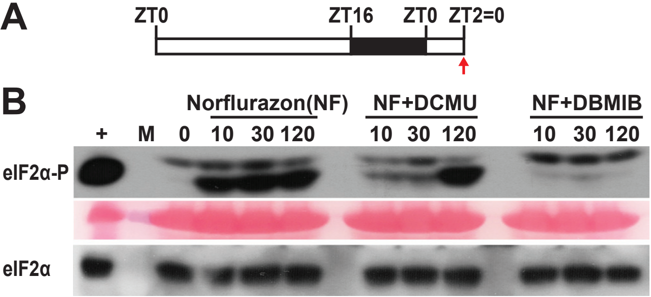


**Figure 11. Norflurazon induces GCN2 activity in a manner dependent on photosynthesis.** Supports Figure 7.
**(A)** Schematic of 16 hr light and 8 hr dark growth regimen and time of norflurazon treatment starting at ZT2 (red arrow). Time=0 represents sampling before treatment.
**(B)** Time course of eIF2$\alpha$ phosphorylation in 14-days-old wild-type Landsberg seedlings. Seedlings were sprayed with either 50μM Norflurazon (NF) or, 30 minutes prior to norflurazon treatment, sprayed with inhibitors of photosynthetic electron transport 8μM DCMU or, 16μM of DBMIB. For details see legend to Supplemental Figure.1.
