## Supplemental Figure 12 for "Light activates the translational regulatory GCN2 kinase via reactive oxygen species emanating from the chloroplast"

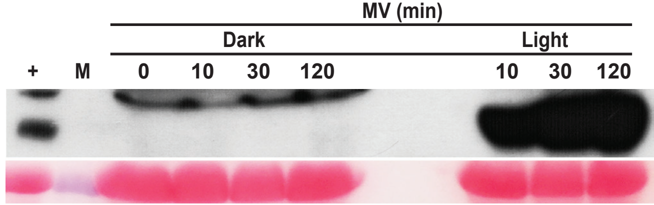


**Supplemental Figure 12. Activation of GCN2 by methyl viologen requires light.** Supports Figure 7.
Time course of eIF2$\alpha$ phosphorylation in 14-days-old wild-type Landsberg seedlings treated with 20µM methyl viologen (MV). Seedlings were either dark acclimated for 24 hr and sprayed with MV (Dark), or without acclimation at ZT2 (Light). For details see legend to Supplemental Figure 1.
