## Supplemental Figure 13 for "Light activates the translational regulatory GCN2 kinase via reactive oxygen species emanating from the chloroplast"

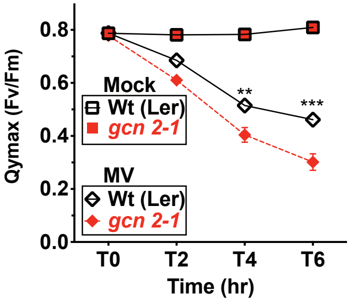


**Supplemental Figure 13 . Effect of methyl viologen on photosynthetic efficiency of wild-type and *gcn2-1*.** Supports Figure 7.
Changes in PSII maximal quantum yield (Fv/Fm) of 16 hr light, 8 hr dark grown rosette stage wild-type Landsberg (Wt) and *gcn2-1* mutant *(gcn2-1)* leaves sprayed with DMSO (Mock) or 20μM methyl viologen (MV) in 0.005% Silwet-77 at ZT4. T0 represents time of first reading after 30 minutes of intial spray. Error bars represent standard error of the mean from fourteen biological replicates. Welch's t-test ***P*-value <0.001, ****P*-value <0.0001 .
