## Supplemental Figure 14 for "Light activates the translational regulatory GCN2 kinase via reactive oxygen species emanating from the chloroplast"

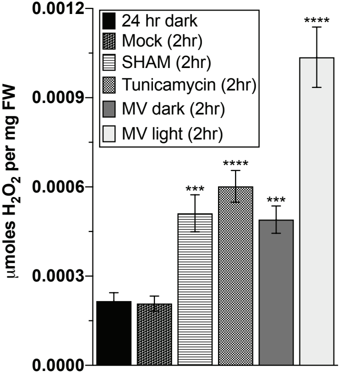


**Supplemental Figure 14. Treatments intended to induce ROS in mitochondria and ER cause accumulation of $H_2O_2$.**
Supports Figure 9.
14-days-old wild-type seedlings were dark adapted for 24 hr (24 hr dark) and then treated with DMSO (Mock), 200μM SHAM, 5μg/ml Tunicamycin for 2 hr. 10μM methyl viologen (MV) was applied either after dark adaptation (MV dark), or at ZT2 in the light (MV light). $H_2O_2$ content was measured with the Amplex red assay. Error bars represent standard error of the mean of 8 biological replicates. Welch's *t*-test *P*-value**** >0.0001.
