## Supplemental Figure 15 for "Light activates the translational regulatory GCN2 kinase via reactive oxygen species emanating from the chloroplast"

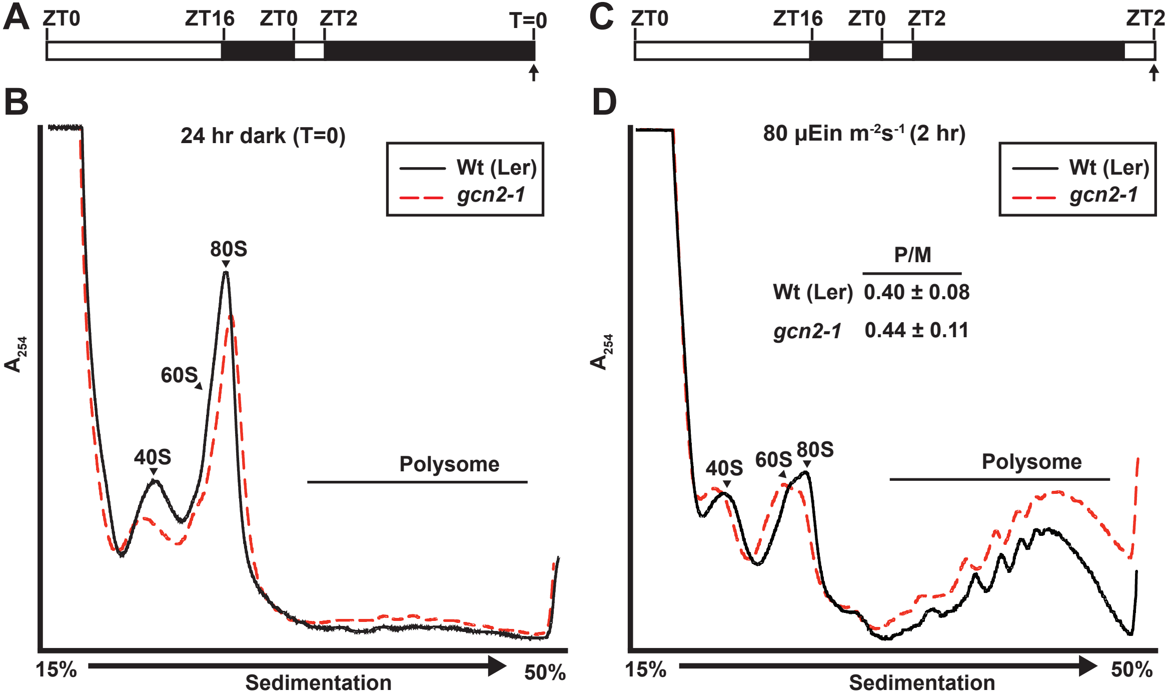


**Supplemental Figure 15. *gcn2* mutant and wild-type show similar ribosome RNA profile upon dark adaptation and after return to light.** Supports Figure 1.
**(A, C)** Schematic of the light treatment. Arrows indicated the time of sampling for panels (B) and (D). (ZT= Zeitgeber time)
Absrobance profile of 14-days-old wild-type Landsberg (Wt(Ler)) and *gcn2-1* mutant *(gcn2-1)* seedlings after 24hr dark acclimation **(B)** and after an additional 24 hr light **(D)**. Cell extracts were fractionated on 15-50% sucrose gradients. Positions of the 40S, 60S, 80S ribosomes and polysomes are indicated. The polysome to monosome (P/M) ratio with standard deviation from five biological replicates is indicated in panel D (Student's *t-test P* >0.5).
