## Supplemental Figure 16 for "Light activates the translational regulatory GCN2 kinase via reactive oxygen species emanating from the chloroplast"

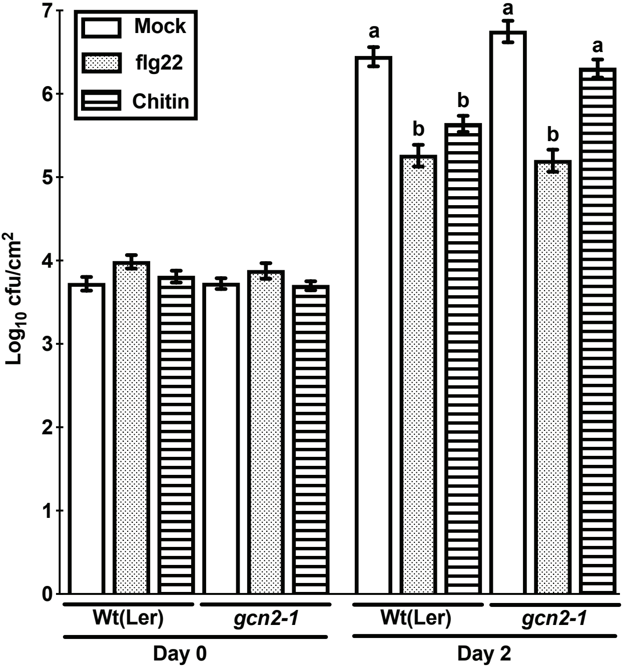


**Supplemental Figure 16. GCN2 imparts bacterial resistance by chitin-inducible priming**. Supports Figure 12.
Leaves of 5-weeks-old wild-type Landsberg Wt(Ler) and *gcn2-1* mutant *(gcn2-1)* were infiltrated with 1µM flg22 (flg22), 1µM chitin (Chitin) or water (Mock) one day before bacterial inoculations. *Pst* DC3000 was infiltrated at a concentration of 2 x $10^5$ cfu $ml^{-1}$ and bacterial growth was enumerated at 2 days post inoculation. Error bars represent standard error of the mean from five independent biological repeats with each having three technical replicates (n =15) (one-way ANOVA, Tukey test; $P < 0.05$).
