## Supplemental Table 1 for "Light activates the translational regulatory GCN2 kinase via reactive oxygen species emanating from the chloroplast"

**Supplemental Table 1. Functional enrichment of mRNA clusters that respond at the translation level to GCN2 or herbicide.** Selected enriched terms are listed with their fold-enrichment over the background set, the FDR-corrected p-value, and the number of genes out of the background set that were discovered in the given cluster. Terms belong to the gene ontology domains biological process (BP), cellular compartment (CC), or molecular function (MF). Related to Figure 10E.

| Gene ontology term | Enrichment | FDR-value | Discovery | Domain |
|---|---|---|---|---|
| **Cluster 1** | | | | |
| ER | 2.1 | 0.0006 | 42 / 716 | CC |
| Vesicle-mediated transport | 3.1 | 0.0006 | 24 / 295 | BP |
| ER-to-Golgi vesicle transport | 7.3 | 0.0018 | 8 / 41 | BP |
| Mitochondrion | | | | CC |
| Mitochondrial envelope | 2.5 | 0.0065 | 20 / 288 | CC |
| Mitochondrial inner membrane | 2.8 | 0.0039 | 18 / 229 | CC |
| NTP biosynthesis | 6.8 | 0.0169 | 6 / 31 | BP |
| H+ transmembrane transport | 6.6 | 0.0015 | 9 / 51 | BP |
| H+ ATPase, rotenone sensitive | 19.2 | 0.0003 | 5 / 10 | MF |
| H+ transporting ATP synthase | 7.6 | 0.0400 | 4 / 19 | CC |
| Other enriched | | | | |
| Histone exchange | 23.1 | 0.0160 | 3 / 5 | BP |
| Gamma-tubulin ring complex | 25.0 | 0.0485 | 2 / 3 | CC |
| Ribosome | 2.4 | 0.0065 | 22 / 348 | CC |
| Depleted | | | | |
| ATP binding | 0.1 | 1.6 10^-10 | 5 /1564 | MF |
| Phosphorylation | 0.1 | 0.0004 | 3 / 849 | BP |
| Regulation of N-compound metabolism | 0.5 | 0.0162 | 22 /1694 | BP |
| Oxoacid metabolism | 0.4 | 0.0158 | 9 / 971 | BP |
| Regulation of gene expression | 0.5 | 0.0024 | 21 /1734 | BP |
| **Cluster 2** | | | | |
| Defense response | | | | |
| Killing cells of other organism | 85.7 | 6.4 10^-6 | 5 / 18 | BP |
| Extracellular region | 9.0 | 3.5 10^-8 | 15 / 571 | CC |
| Peptidase inhibitor | 62.5 | 0.0014 | 4 / 23 | MF |
| Depleted | | | | |
| Nucleus | 0.0 | 0.0008 | 0 / 2923 | CC |
| **Cluster 3** | | | | |
| Protein storage vacuole | 60.0 | 0.0075 | 3 / 10 | CC |
