## Supplemental Table 2 for "Light activates the translational regulatory GCN2 kinase via reactive oxygen species emanating from the chloroplast"

**Supplemental Table 2. Upon activation of GCN2 by herbicide, the resultant global loss of ribosome loading spares certain functional categories of mRNA from translational repression.** For this analysis, the raw mRNA translation states were adjusted to acknowledge that herbicide triggers a global repression of ribosome loading in the wild type. This was done by reducing the translation states of wild type treated with chlorosulfuron herbicide by the global drop in ribosome loading, averaged from the biological replicates. Translationally repressed mRNAs were then identified by limma (3,694 mRNAs) and analyzed for enriched (repressed) and depleted (spared) terms by TopGO/Panther. Gene ontology terms are listed with their fold-enrichment over the background set, the FDR-corrected p-value, and the number of genes out of the background set that were discovered as being translationally repressed. Terms belong to the gene ontology domains biological process (BP), cellular compartment (CC), or molecular function (MF).

| Gene ontology term | Enrichment | FDR-value | Discovery | Domain |
|---|---|---|---|---|
| **Translationally repressed** | | | | |
| Translation | 2.2 | 6 10^ -50 | 311 / 601 | BP |
| Ribosome | 3.5 | 1 10^-115 | 282 / 348 | CC |
| Nitrogen | | | | |
| Amide transmembrane transport | 2.3 | 4.3 10^ -5 | 31 / 59 | MF |
| Peptidase, Thr-type, endo- | 2.9 | 2.2 10^ -4 | 16 / 24 | MF |
| Proteasome core | 3.0 | 5.2 10^ -5 | 16 / 23 | CC |
| Autophagosome membrane | 3.5 | 8.5 10^ -4 | 9 / 11 | CC |
| Mitochondrion | | | | |
| Inner membrane | 2.3 | 3.7 10^ -22 | 124 / 229 | CC |
| Envelope | 2.1 | 2.0 10^ -20 | 142 / 288 | CC |
| Chloroplast | | | | |
| Thylakoid | 1.7 | 4.1 10^ -13 | 158 / 387 | CC |
| Nucleus | | | | |
| Nucleosome | 3.3 | 6.8 10^ -11 | 29 / 37 | CC |
| RNA polymerase | 1.9 | 1.4 10^ -2 | 25 / 58 | MF |
| **Spared from translational repression** | | | | |
| Translation | | | | |
| tRNA aminoacylation | 0.10 | 5.3 10^ -3 | 1 / 44 | BP |
| Nitrogen | | | | |
| Cellular amino acid metabolism | 0.38 | 3.7 10^ -11 | 32 / 364 | BP |
| Nitrate assimilation | 0.25 | 1.4 10^ -1 | 2 / 34 | BP |
| Amino acid transport | 0.16 | 3.8 10^ -3 | 2 / 54 | BP |
| Alpha-amino acid catabolism | 0.34 | 3.9 10^ -2 | 5 / 62 | BP |
| Cell division | 0.65 | 1.1 10^ -2 | 45 / 297 | BP |
| Carbohydrate metabolism | 0.41 | 1.2 10^ -20 | 72 / 747 | BP |
| Starch metabolism | 0 | 3.2 10^ -5 | 0 / 56 | BP |
| Response to stress | | | | |
| Defense response, general | 0.67 | 2.4 10^ -7 | 145 / 924 | BP |
| Ethylene metabolism | 0 | 2.9 10^ -2 | 0 / 25 | BP |
| Phenylpropanoid metabolism | 0.37 | 1.2 10^ -3 | 10 / 117 | BP |
| Cyclic nucleotide binding | 0 | 3.9 10^ -2 | 0 / 24 | MF |
| Hydrogen peroxide metabolism | 0.40 | 9.2 10^ -2 | 6 / 64 | BP |
| Oxylipin biosynthesis | 0.20 | 3.2 10^ -1 | 1 / 21 | BP |
| Protein kinase | 0.26 | 1.4 10^ -31 | 39 / 654 | MF |
| Nucleus | | | | |
| Gene silencing | 0.34 | 1.2 10^ -4 | 11 / 139 | BP |
| Helicase | 0.16 | 1.5 10^ -9 | 6 / 158 | MF |
